# Foxp4^+^ Mandibular Skeletal Stem Cells Orchestrate Bone/Tooth Development and Regeneration

**DOI:** 10.64898/2026.04.29.721551

**Authors:** Lei Zhang, Dandan Cao, Xinyu Li, Hongqiang Yu, Jianfang Wang, Qiaoling Zhu, Xinyi Zhang, Chen Chen, Gongchen Li, Xinyu Xu, Xiaoqiao Xu, Dike Tao, Xuyan Gong, Pingping Niu, Xiaoshan Wu, Mengfei Yu, Rui Yue, Yao Sun

**Affiliations:** Shanghai Engineering Research Center of Tooth Restoration and Regeneration & Tongji Research Institute of Stomatology & Department of Implantology, Shanghai Tongji Stomatological Hospital and Dental School, Tongji University, Shanghai 200072, China; State Key Laboratory of Cardiovascular Diseases and Medical Innovation Center, Shanghai East Hospital, Frontier Science Center for Stem Cell Research, Shanghai Key Laboratory of Signaling and Disease Research, School of Life Sciences and Technology, Tongji University, Shanghai 200092, China; Shanghai Academy of Natural Sciences (SANS), Tongji University, Shanghai 200092, China; Shanghai Institute of Stem Cell Research and Clinical Translation, Shanghai 200120, China; Shanghai Engineering Research Center of Tooth Restoration and Regeneration & Tongji Research Institute of Stomatology & Department of Oral and Maxillofacial Surgery, Shanghai Tongji Stomatological Hospital and Dental School, Tongji University, Shanghai 200072, China; Stomatology Hospital, School of Stomatology, Zhejiang University School of Medicine, Zhejiang Provincial Clinical Research Centefor Oral Diseases, Zheiiang Key Laboratory of Oral Biomedical, Hangzhou 310000, China; Academician Workstation for Oral-Maxillofacial Regenerative Medicine, Central South University, Changsha 410013, China; Department of Oral and Maxillofacial Surgery, Xiangya Hospital, Central South University, Changsha 410028, China

**Author notes:** These authors contributed equally (Lei Zhang, Dandan Cao, Xinyu Li, Hongqiang Yu). Correspondence (R.Y.), (Y. S.).

**Keywords:** Mandible development, Skeletal stem cells, Primary cilia, Tooth regeneration, Distraction osteogenesis

## Abstract

The identity of stem/progenitor cells that drive mandible/tooth development and regeneration remains poorly understood. Here, we show that cranial neural crest-derived Foxp4^+^ mesenchymal cells condense to form Meckel’s cartilage, and simultaneously promote mandible development and tooth germ formation. In postnatal mandibles, Foxp4^+^ cells localize in the endosteum, periosteum and dental pulp, which undergo osteogenesis and odontogenesis under steady state, as well as chondrogenesis during fracture repair. Importantly, transplantation of embryonic or adult Foxp4^+^CD200^+^CD105^-^ mandibular skeletal stem cells (mdSSCs) generate ectopic bone and tooth. In contrast, genetic ablation of Foxp4^+^ cells severely impairs mandible/tooth development and delays fracture healing. Mechanistically, we find that primary cilia critically regulate mdSSC maintenance and differentiation into bone and tooth. FOXP4^+^PDPN^+^CADM1^+^ mdSSCs also exist in adult human mandible, which generate ectopic bone and intact tooth upon transplantation into immunodeficient mice. Taken together, we identify Foxp4^+^ mdSSCs with potential therapeutic applications in regenerating mandibular bone and tooth.

## Introduction

Micrognathia is a severe and prevalent craniofacial abnormality, which mainly affects the mandible.^1–5^ Surgical interventions such as distraction osteogenesis (DO) remain the primary treatments for micrognathia,^6–8^ which are often accompanied by infection, nonunion/malunion and prolonged rehabilitation period during treatment.^9,10^ Efficient repair of the mandibular bones necessitates deeper understanding of their developmental mechanisms.^11,12^ In contrast to calvarial bones that are formed by direct ossification of the osteoprogenitors,^13^ the mandibles are developed from Meckel’s cartilage (MC), a unique cartilaginous tissue that serves as the embryonic template for mandible and middle ear development.^12,14–16^ MC is generated by chondrogenic differentiation of cranial neural crest-derived cells (NCDCs) to promote elongation of the embryonic mandible,^12,17^ which is then ossified by osteoprogenitors and quickly disappears after birth.^18,19^ However, the identity of NCDCs that specify MC formation to initiate mandible development, as well as the stem/progenitor cells governing maintenance and regeneration of mandibles in the adulthood remain poorly understood.

Craniofacial abnormalities are often accompanied by tooth dysplasia and anodontia.^20–22^ Teeth are highly specialized hard tissues indispensable for food intake, language communication and facial aesthetics.^23^ Tooth loss induced by trauma, periodontal diseases and aging can lead to malnutrition and compromised life quality.^24^ Current treatments for tooth lost predominantly rely on dental implantation, which faces multiple challenges such as poor osseointegration and alveolar bone resorption.^25^ Up till now, autologous tooth regeneration has not been achieved and remains an unmet medical need. Tooth development initiates with tooth germ formation involving close interaction between epithelial and mesenchymal cells, which generate enamel and dentin/dental pulp, respectively.^26^ However, the reason why teeth are only found in maxilla and mandible, as opposed to other parts of the skeleton, has not been uncovered. Interestingly, the mesenchymal cells forming tooth germ, MC and mandibular bone are all cranial NCDCs,^27^ whose developmental defect could lead to craniofacial and tooth abnormalities at the same time.^12,28,29^ Whether a multipotent stem/progenitor population exists in the cranial NCDCs to generate tooth, MC and mandibular bone during early development remain to be explored. Furthermore, whether stem/progenitor cells capable of regenerating bone/tooth retain in the adult mandible are still unknown.

Skeletal development and remodeling are orchestrated by region-specific skeletal stem cells (SSCs).^30–32^ SSCs were initially identified and extensively studied in the bone marrow stroma.^33–36^ Mouse and human SSCs were also found in the growth plate, which critically regulate endochondral ossification and elongation of the long bones.^37–39^ Periosteal SSCs promote osteogenic differentiation by intramembranous ossification under steady state, and acquire chondrogenic capacity upon injuries to repair bone fractures.^40,41^ Apart from the appendicular skeleton, recent studies discovered vertebral and calvarial SSCs in the axial skeleton that not only regulate skeletogenesis, but also underlie both tumor metastasis and craniosynostosis.^42,43^ Our previous study identified FOXP1/2/4^+^ embryonic skeletal stem/progenitor cells (eSSPCs) in the perichondrium of 8-week-old human embryonic long bones and calvaria.^44^ However, whether eSSPCs contribute to mandible and/or tooth development remains unknown. In addition to SSCs, previous studies showed that dental stem/progenitor cells, including Gli1^+^ cells,^45^ NG2^+^ cells ^46^ and Plp1^+^ cells^47^ mediate incisor development and repair. In contrast, the identity of stem/progenitor cells indispensable for molar development and regeneration have been elusive.

In this study, we reported the discovery of Foxp4^+^ mandibular SSCs (mdSSCs), a pivotal subset of cranial NCDCs that tightly regulates the development and regeneration of mandibular bones and molars.

## Results

### Embryonic Foxp4^+^ cells give rise to MC, mandibular bone and molar tooth germ mesenchyme

Mandible development starts from embryonic day 12.5 (E12.5), when cranial NCDCs condense to initiate MC formation. From E15.5 onward, mineralized bone tissue begins to encapsulate MC, followed by cartilage resorption (Figure S1A). To identify skeletal stem/progenitor cells governing mandible development, we analyzed our published single-cell RNA-sequencing (scRNA-seq) dataset of 8-week-old human embryonic calvaria.^44^ Compared to eSSPC markers *FOXP1 and FOXP2*,^44^ *FOXP4* is more specifically expressed in NCDCs and migratory neural crest cells (Figure 1A). We also performed immunofluorescence (IF) staining with the endogenous antibodies against Foxp1/2/4, and confirmed that Foxp4 exhibits a more restricted expression pattern in E15.5 mandible as compared to Foxp1/2 (Figures S1B-S1D). Therefore, we constructed a *Foxp4-CreERT2* knock-in allele and crossed it with *Rosa26-tdTomato* reporter mice (*Foxp4-CreERT2; tdTomato* hereafter) to perform genetic lineage tracing of Foxp4^+^ cells at different developmental stages (Figure 1B).

**Figure 1.**
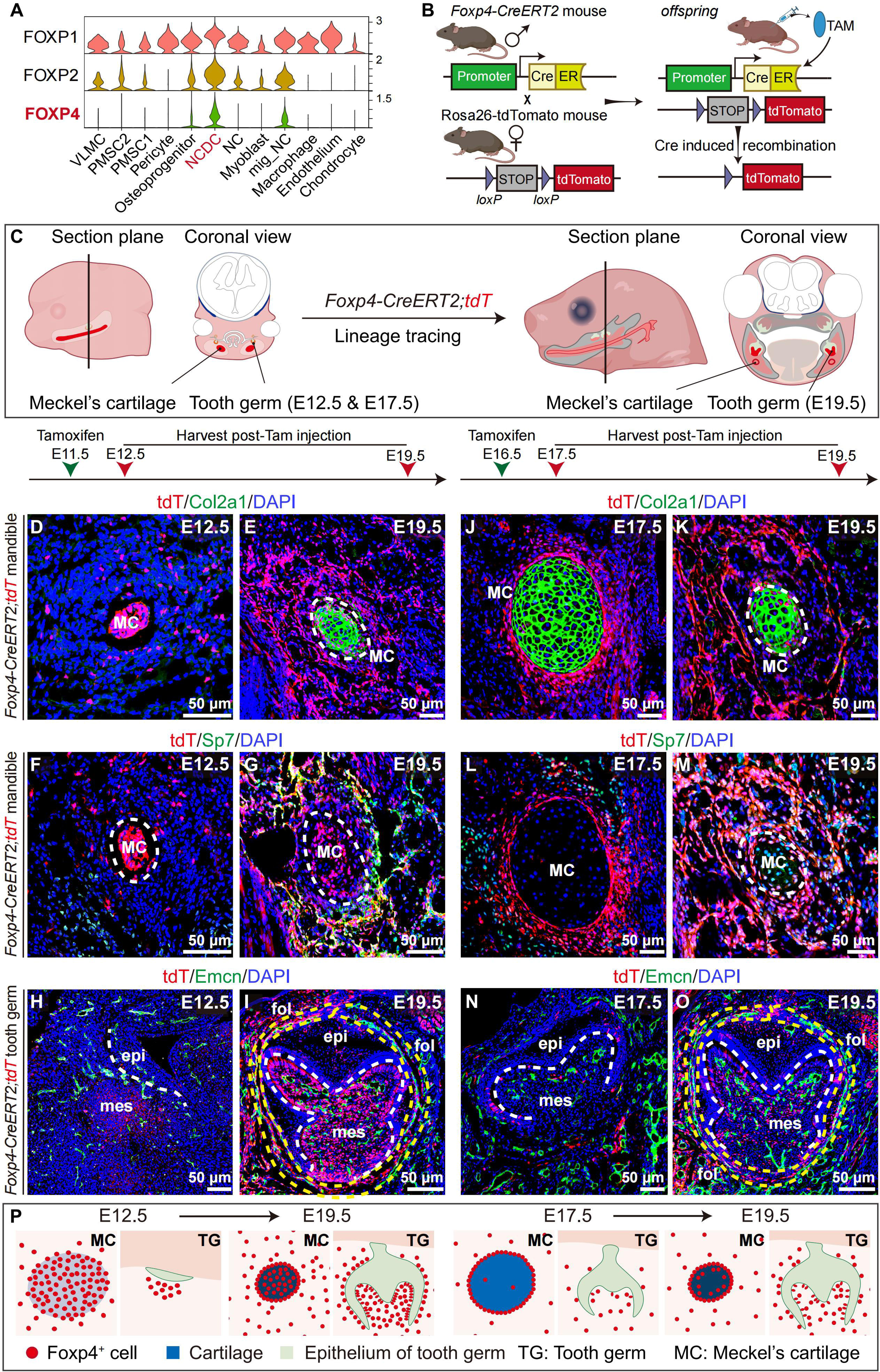
Foxp4^+^ cells contribute to mandible and tooth development. (A) Violin plots showing the expression pattern of FOXP1/2/4 genes across different cell clusters in our previously published 8-week-old human embryonic calvarial scRNA-seq data (GSE143753). (B) Schematic illustration of the genetic lineage tracing strategy. (C) Lineage tracing strategy of Foxp4^+^ cell during mandible and tooth development. (D-G) IF images of E12.5 and E19.5 mandibles in *Foxp4-CreERT2; tdTomato* mice. Tamoxifen was administered at E11.5. The tdTomato (tdT), Col2a1, Sp7 and DAPI signals were shown. The dotted circles indicated Meckel’s cartilage (MC) (n=3 mice per group from 3 independent experiments). (H and I) IF images of E12.5 and E19.5 tooth germs in *Foxp4-CreERT2; tdTomato* mice. Tamoxifen was administered at E11.5. The tdTomato (tdT), Emcn (blood vessel marker) and DAPI signals were shown. White dotted lines indicated the border between odontogenic epithelium and tooth germ mesenchyme. Yellow dotted lines indicated the dental follicle. mes: mesenchyme, epi: epithelium, fol: follicle (n=3 mice per group from 3 independent experiments). (J-M) IF images of E17.5 and E19.5 mandibles in *Foxp4-CreERT2; tdTomato* mice. Tamoxifen was administered at E16.5. The tdTomato (tdT), Col2a1, Sp7 and DAPI signals were shown. The dotted circles indicated Meckel’s cartilage (MC) (n=3 mice per group from 3 independent experiments). (N and O) IF images of E16.5 and E19.5 tooth germs in *Foxp4-CreERT2; tdTomato* mice. Tamoxifen was administered at E16.5. The tdTomato (tdT), Emcn (blood vessel marker) and DAPI signals were shown. White dotted lines indicated the border between odontogenic epithelium and tooth germ mesenchyme. Yellow dotted lines indicated the dental follicle. mes: mesenchyme, epi: epithelium, fol: follicle (n=3 mice per group from 3 independent experiments). (P) Schematic illustration of the lineage tracing results (D-O).

*Foxp4-CreERT2; tdTomato* mice were grossly normal, with similar body size and mandible/tooth development as *Rosa26-tdTomato* littermate controls (Figures S1E-S1G). To test whether Foxp4^+^ cells contribute to MC/tooth germ formation, we administered tamoxifen at E11.5 or E16.5 to induce *Foxp4-CreERT2* recombination (Figures 1C). IF staining of Foxp4 showed co-localization with tdTomato at both E12.5 (data not shown) and E17.5 (Figure S1H), suggesting that *Foxp4-CreERT2* recombination faithfully reflects endogenous Foxp4 expression.

At E12.5, Foxp4^+^ cells condensed to specify MC formation, which generated Col2a1^+^ chondrocytes and Sp7^+^ osteoprogenitors/osteoblasts surrounding MC at E19.5 (Figures 1D-1G). Interestingly, Foxp4^+^ cells also condensed to form the molar tooth germ mesenchyme at E12.5, which generated odontoblasts and dental follicle at E19.5 (Figures 1H and 1I). To further confirm this, we performed tissue-clearing and light-sheet microscopy to trace Foxp4-lineage cells at E19.5 (tamoxifen administered at E11.5), and found that tdTomato^+^ cells were readily detected in the molar tooth germ mesenchyme, as well as the dental follicle (Figures S1K). In contrast, very few Foxp4^+^ cells were found in the maxilla at E12.5 (Figure S1I), with much less contribution to molar tooth germ mesenchyme as compared to their mandibular counterparts (Figure S1J).

At E17.5, Foxp4^+^ cells were found in the perichondrial region of hypertrophic MC, which generated less Col2a1^+^ chondrocyte but abundant Sp7^+^ osteoprogenitors/osteoblasts at E19.5 (Figures 1J-1M). In contrast to E12.5, very few Foxp4^+^ cells localized in molar tooth germ mesenchyme at E17.5, with limited contribution to E19.5 odontoblasts and dental follicle (Figures 1N and 1O). Taken together, these data showed significant contribution of embryonic Foxp4^+^ cells to mandible development and molar tooth germ formation (Figure 1P).

### Identification and functional characterization of Foxp4^+^ mdSSCs in embryonic mouse mandible

Next, we performed scRNA-seq to dissect the cellular heterogeneity within Foxp4^+^ cells. To do this, mandibles were harvested from *Foxp4-CreERT2; tdTomato* embryos at E12.5 (tamoxifen administered at E11.5). CD45^-^ Ter119^-^ CD31^-^ Thy1^-^ tdTomato^+^ cells (non-hematopoietic, non-endothelial, non-fibroblastic Foxp4^+^ cells) were flow cytometrically sorted after enzymatic dissociation, followed by 3’ scRNA-seq on the 10X Genomics platform (Figure 2A). After stringent quality control and doublets removal, we obtained 12,584 single cells in total. Clustering analysis by uniform manifold approximation and projection (UMAP) revealed 7 cell clusters (Figure 2B), all of which expressed the mesenchymal marker *Prrx1* (Figure 2C). Interestingly, cluster 1 expressed the highest level of NCDC marker *Meis2* (Figure 2C),^48^ and represented 29% of all single cells (Figure 2D). Pseudotime analysis by Monocle3 revealed that cluster 1 underwent chondrogenesis, osteogenesis and odontogenesis (Figure 2E).^49^ Cell-cell interaction analysis showed that cluster 1 functions as a signaling hub among all clusters (Figure 2F). We also performed gene regulatory network (regulon) analysis by SCENIC (Figure 2G)^50^ and found that *Meis1*^51^ regulon was most active in cluster 1, while *Sox9*,^52^ *Msx2*^53^ and *Lhx8* ^54^ regulons were highly enriched in the chondrogenic (clusters 5 and 6), osteogenic (cluster 3) and odontogenic (cluster 4) clusters, respectively (Figure 2H). Importantly, we found that cluster 1 expressed mSSC marker *CD200* but was negative for *Eng* (*CD105*) and *Enpep* (*6C3*) (Figure 2C),^37^ suggesting a previously uncharacterized mandibular SSC (mdSSC) population.

**Figure 2.**
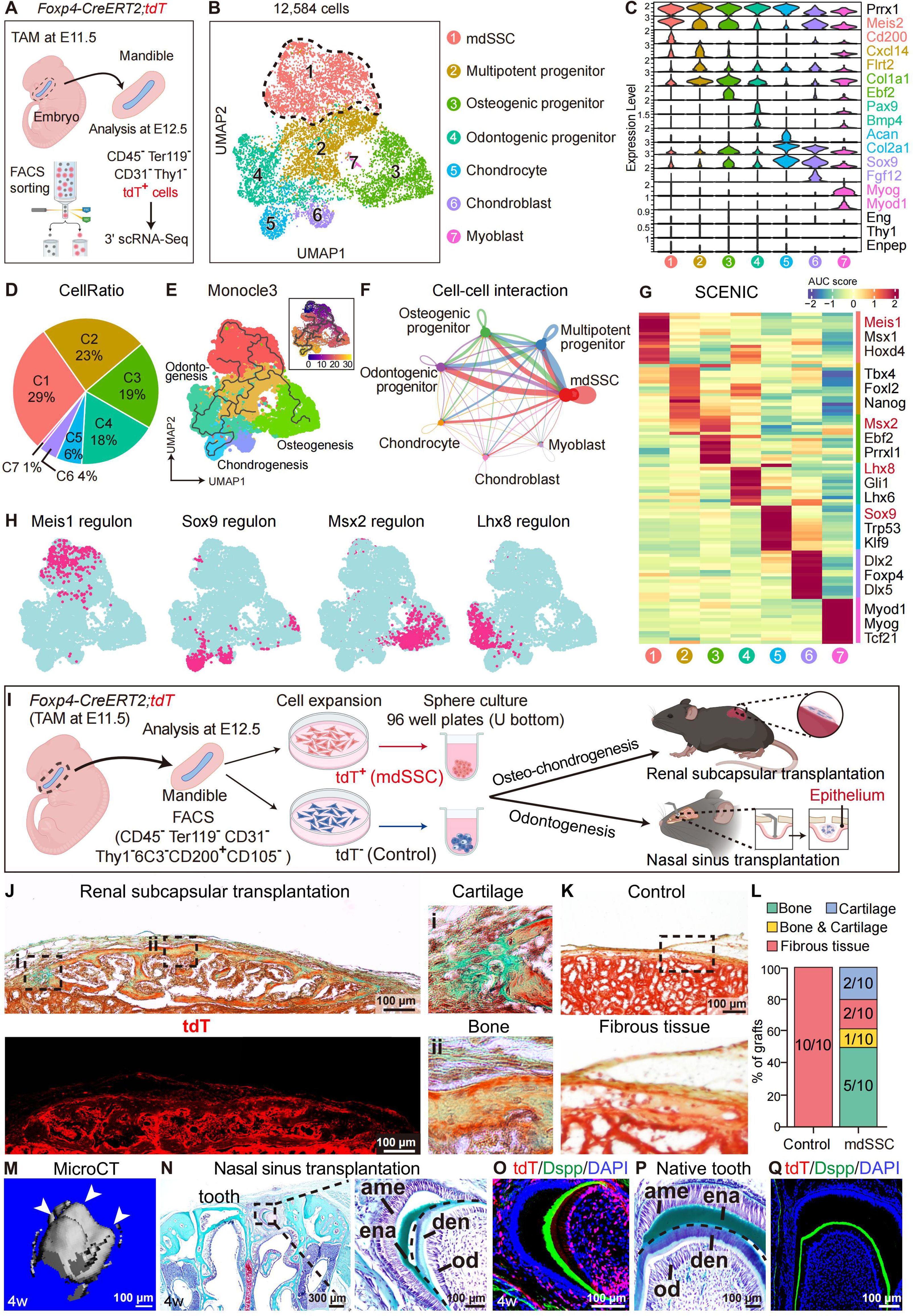
Identification and functional characterization of Foxp4^+^ mdSSCs in E12.5 mouse mandible. (A) scRNA-seq workflow and experimental scheme. (B) UMAP visualization of 7 cell clusters in E12.5 mouse mandible. In total, 12,584 single cells from 5 embryos were analyzed. Dotted lines indicated the mdSSC cluster. (C) Violin plots showing the expression of feature genes across the 7 cell clusters. Marker genes for annotating each cell cluster were color-coded on the right. (D) Pie chart showing the cell ratios of each cell cluster. (E) UMAP visualization of the osteogenic, chondrogenic and odontogenic trajectories simulated by Monocle3. Diffusion pseudotime was indicated in the upper right panel. (F) Circle plot showing cell-cell interactions among the 7 cell clusters. The interaction strength (indicated by line thickness) was calculated by CellChat. (G) Heatmap showing regulons enriched in each cell cluster. The AUC score (row scaling) was computed. Three representative regulons (out of top 10 regulons) for each cell cluster were listed on the right. (H) Binary activities of Meis1, Sox9, Msx2 and Lhx8 regulons were shown by UMAP plots. (I) Transplantation workflow and experimental scheme. (J and K) Movat’s pentachrome staining after renal subcapsular transplantation of Foxp4^+^ mdSSCs (J) or control (K) cells for 4 weeks. Cartilage (i) and bone (ii) formation were shown. The tdTomato (tdT) signal indicated donor-derived cells (J, lower left panel). (L) Quantification of the renal subcapsular transplantation data (n=10 mice per group from 5 independent experiments). (M) Representative microCT image of the ectopic tooth formed by Foxp4^+^ mdSSCs 4 weeks after nasal sinus transplantation. Arrowheads indicated the tooth cups. (N and O) Safranin O/Fast Green (N) and IF (O) staining of the ectopic tooth formed by Foxp4^+^ mdSSCs 4 weeks after nasal sinus transplantation. Foxp4^+^ mdSSCs from E12.5 *Foxp4-CreERT2; tdTomato* embryos were transplanted. ame: ameloblast, ena: enamel, den: dentin, od: odontoblast. (P and Q) Safranin O/Fast Green (P) and IF (Q) staining of the native tooth in the recipient mice. ame: ameloblast, ena: enamel, den: dentin, od: odontoblast.

To functionally test the stem cell activity of Foxp4^+^ mdSSCs, we flow cytometrically sorted CD45^-^ Ter119^-^ CD31^-^ Thy1^-^ 6C3^-^ CD200^+^ CD105^-^ cells from E12.5 *Foxp4-CreERT2; tdTomato* mandibles (tamoxifen administered at E11.5), and sub-divided them into tdTomato^+^ (mdSSCs) or tdTomato^-^ (control) fractions (Figures S2A and S2B). Colony-forming unit-fibroblast (CFU-F) assay showed significantly higher clonogenic activity of mdSSCs (Figure S2C). *In vitro* trilineage differentiation analyses showed significantly increased osteogenic, adipogenic and chondrogenic capacities in mdSSCs as compared to control cells (Figures S2D-S2F). To test their osteo-chondrogenic potential *in vivo*, we performed renal subcapsular transplantation of mdSSCs or control cells after 2D expansion and sphere culture (Figure 2I). Four weeks after transplantation, mdSSCs gave rise to ectopic bone and cartilage tissues efficiently (Figures 2J and 2L), while control cells mainly generated fibrous tissue (Figures 2K and 2L).

Considering that tooth development requires close interaction between epithelial and mesenchymal cells,^23,55,56^ and that the nasal sinus epithelium is functionally similar to odontogenic epithelium,^57–59^ we established a nasal sinus transplantation model where mdSSCs or control cells were implanted on top of the nasal sinus epithelium after 2D expansion and sphere culture (Figure 2I). MicroCT analysis revealed that mdSSCs, but not control cells, generated ectopic tooth with multiple cusps 4 weeks after nasal sinus transplantation (Figure 2M). Safranin O/Fast Green staining showed dental pulp, as well as ameloblasts and odontoblasts that formed enamel and dentin, respectively (Figures 2N and 2O), resembling the native mouse tooth (Figures 2P and 2Q). Nasal sinus transplantation of mdSSCs also generated ectopic bone and cartilage as revealed by histological analyses, while control cells mainly generated fibrous tissue (Figures S2G-S2L). We also performed nasal sinus transplantation of E15.5-E16.5 mdSSCs from mouse mandibles, and found that they retain the ability to regenerate ectopic tooth (containing dental crown with cusps and dental pulp) 4 weeks after transplantation (Figures S2M-S2O). Remarkably, we observed evident dental root formation with root apex 8 weeks after transplantation (Figure S2P), suggesting continued maturation of the ectopic tooth.

Taken together, we identified Foxp4^+^ mdSSCs in embryonic mouse mandible that can be prospectively isolated and give rise to ectopic bone, cartilage and tooth upon *in vivo* transplantation.

### Postnatal Foxp4^+^ cells undergo osteogenesis and odontogenesis at steady state, as well as chondrogenesis during fracture repair

To analyze the fate of Foxp4^+^ cells in postnatal mice, we administered tamoxifen in neonatal (P1 to P3) or adult (12-week-old) *Foxp4-CreERT2; tdTomato* mice to perform lineage tracing, and analyze their distribution in the mandibular bone and M1 molar (Figures S3A and S3B). At P4, Foxp4^+^ cells localize in the endosteal, periosteal and mesenchymal regions of the nascent mandibular bone (Figures 3A). Foxp4^+^ cells were also detected in the dental pulp and periodontal membrane (Figure 3B). After 12 weeks of lineage tracing, Foxp4^+^ cells gave rise to abundant endosteal/periosteal cells, bone marrow perivascular stromal cells, and osteocytes in the mandibular bone (Figure 3C), as well as molar odontoblasts, periodontal tissues, alveolar bone lining cells and osteocytes (Figure 3D). When we administered tamoxifen in 12-week-old *Foxp4-CreERT2; tdTomato* mice, more Foxp4^+^ cells were detected in the endosteal as compared to periosteal regions in the mandibular bone (Figure 3E). Foxp4^+^ cells were also detected in the pulp cavity of dental root and periodontal regions at this stage (Figure 3F). After 12 weeks of lineage tracing, Foxp4^+^ cells gave rise to abundant endosteal/periosteal cells, bone marrow perivascular stromal cells and osteocytes in 24-week-old mandibular bone (Figure 3G). Foxp4^+^ cells also generated most odontoblasts in the dental root, as well as a large proportion of periodontal tissues, alveolar bong lining cells and osteocytes in 24-week-old molar (Figure 3H). Co-staining with Ctsk,^60,61^ Postn^62^ or Dspp^63^ was performed to confirm the contribution of Foxp4^+^ cells to endosteum/periosteum, periodontal ligament and odontoblasts, respectively, after lineage tracing (Figures S3C-S3H).

**Figure 3.**
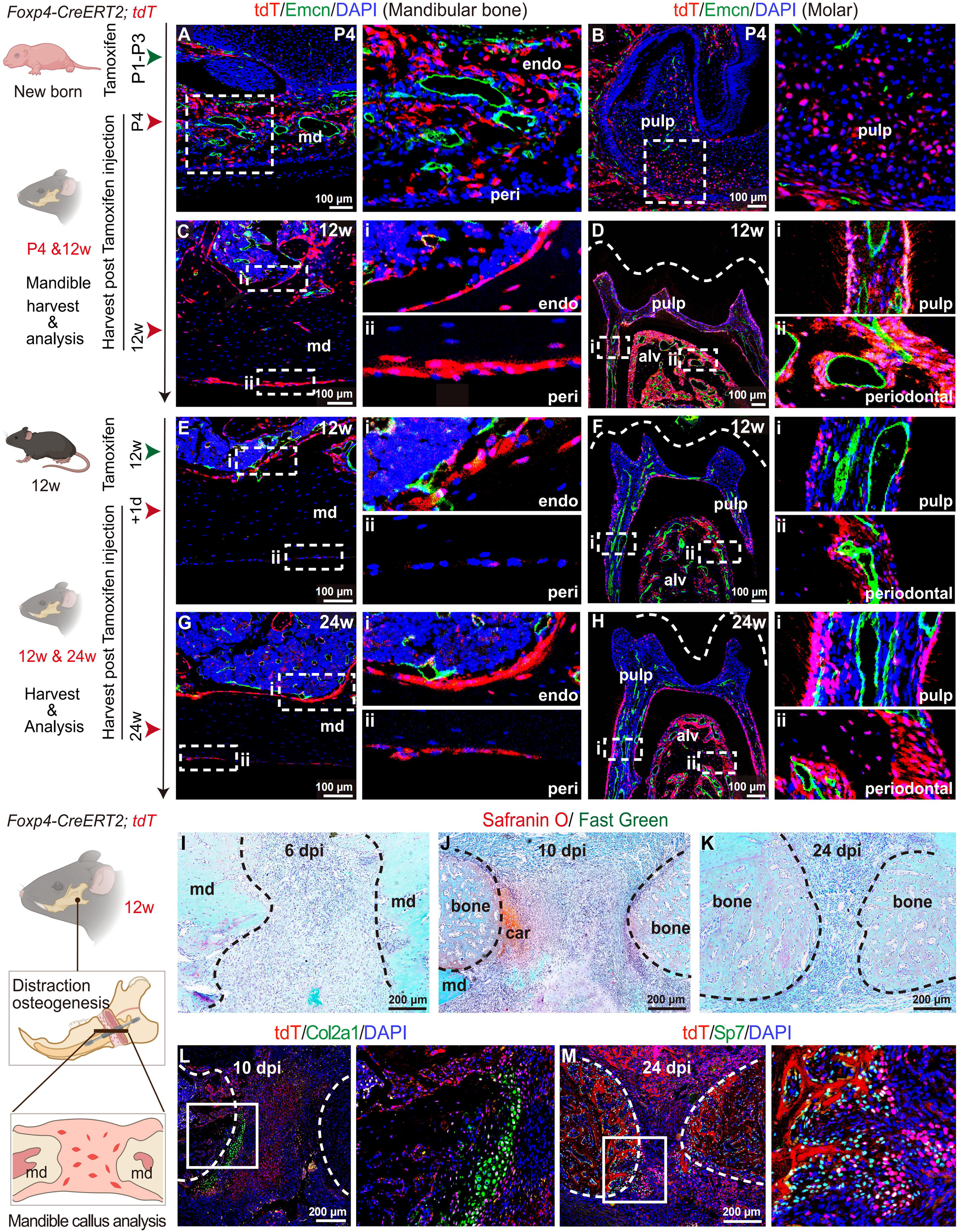
Foxp4^+^ cells undergo osteogenesis and odontogenesis during postnatal mandible development, as well as chondrogenesis during fracture repair in adulthood. (A-D) IF images of P4 and 12-week-old mandible from *Foxp4-CreERT2; tdTomato* mice. The endosteal (endo, i) and periosteal (peri, ii) regions of the mandibular bones (A and C), as well as the dental pulp (pulp, i) and periodontal regions (periodontal, ii) of the molar tooth (B and D) were enlarged. Tamoxifen was administered for 3 consecutive days from P1 to P3. Mice were analyzed after 1 day (A and B) or 12 weeks (C and D). The tdTomato (tdT), Emcn and DAPI signals were shown. Dotted lines (D) indicated the border of the dental crown (n=3 mice per group from 3 independent experiments). md: mandibular bone; alv: alveolar bone. (E-H) IF images of 12- and 24-week-old mandible from *Foxp4-CreERT2; tdTomato* mice. The endosteal (endo, i) and periosteal (peri, ii) regions of the mandibular bones (E and G), as well as the dental pulp (pulp, i) and periodontal regions (periodontal, ii) of the molar tooth (F and H) were enlarged. Tamoxifen was administered for 5 consecutive days starting at 12 weeks of age. Mice were analyzed after 1 day (E and F) or 12 weeks (G and H). The tdTomato (tdT), Emcn and DAPI signals were shown. Dotted lines (F and H) indicated the border of the dental crown (n=3 mice per group from 3 independent experiments). md: mandibular bone; alv: alveolar bone. (I-K) Safranin O/Fast Green staining of mandibular bone repair. DO-operated 12-week-old *Foxp4-CreERT2; tdTomato* mice were analyzed at 6, 10, 24 dpi. Dotted lines indicated the osteogenic front. md: mandible, bone: newly formed woven bone, car: newly formed cartilage (n = 3 mice per group from 3 independent experiments). (L and M) IF images of 10 and 24 dpi mandibular bone after DO surgery. Tamoxifen was administered for 5 consecutive days in 12-week-old *Foxp4-CreERT2; tdTomato* mice before DO surgery. The tdTomato (tdT), Col2a1 (L), Sp7 (M) and DAPI signals were shown (n = 3 mice per group from 3 independent experiments).

To test whether Foxp4^+^ cells undergo chondrogenesis to repair mandible fractures, we established a mouse model of distraction osteogenesis (DO) in 12-week-old *Foxp4-CreERT2; tdTomato* mice,^7^ which mimics the surgical procedure for treating mandibular malformations.^64–66^ Safranin O/Fast green staining showed cartilage formation in the bone callus 10 days post injury (dpi), which was gradually absorbed and mineralized into mature mandibular bone from 17 to 45 dpi (Figures 3I-3K and S3I-S3K). Consistent with this, IF staining showed that Foxp4^+^ cells gave rise to Col2a1^+^ chondrocytes in the bone callus at 10 dpi (Figure 3L), as well as Sp7^+^ osteoprogenitors and osteocytes from 17 to 45 dpi (Figures 3M and S3L-S3N). Taken together, we found that Foxp4^+^ cells undergo osteogenesis and odontogenesis during postnatal mandible growth, and transiently generate chondrocytes to promote callus formation and fracture repair in adult mice.

### Identification and functional characterization of Foxp4^+^ mdSSCs in adult mouse mandible

To test whether mdSSCs exist in adult mice mandible, we harvested molar-extracted mandible from 12-week-old *Foxp4-CreERT2; tdTomato* mice 2 days after tamoxifen administration (5 consecutive days). CD45^-^ Ter119^-^ CD31^-^ Thy1^-^ tdTomato^+^ cells were flow cytometrically sorted after bone crushing and enzymatic dissociation, followed by 3’ scRNA-seq on the 10X Genomics platform (Figure 4A). After stringent quality control and doublets removal, we obtained 11,862 single cells in total. Clustering analysis by UMAP revealed 5 cell clusters (Figure 4B), most of which expressed mesenchymal marker *Prrx1* except cluster 4 (Figure 4C). Since cluster 4 highly expressed epithelial markers *Trp63* and *Krt5*,^67–69^ it might represent oral epithelial cells. Importantly, we found that cluster 1 (22% of all single cells, Figure 4D) highly expressed both chondrocyte (*Acan, Col2a1*) and osteoblast (*Col1a1, Pth1r*) marker genes (Figure 4C). Cluster 1 also expressed mSSC marker *CD200* but was negative for *Eng* (*CD105*) and *Enpep* (*6C3*) (Figure 4C), suggesting an adult mdSSC population. Pseudotime analysis by Monocle3 revealed that cluster 1 underwent osteogenesis to generate osteoprogenitors (cluster 3), and fibrogenesis to generate periosteal/periodontal cells (cluster 5) (Figure 4E). Cell-cell interaction analysis revealed that cluster 1 functioned as a signaling hub among different cell clusters (Figure 4F). Gene regulatory network analysis showed that the Sox9 regulon was highly enriched in cluster 1, while the Sp7 regulon became increasingly actively along the osteogenic differentiation trajectory (Figures 4G and 4H). In contrast, the Npdc1^70^ and Trp63 regulons were enriched in the periosteal/periodontal and epithelial cell clusters, respectively (Figures 4G and 4H).

**Figure 4.**
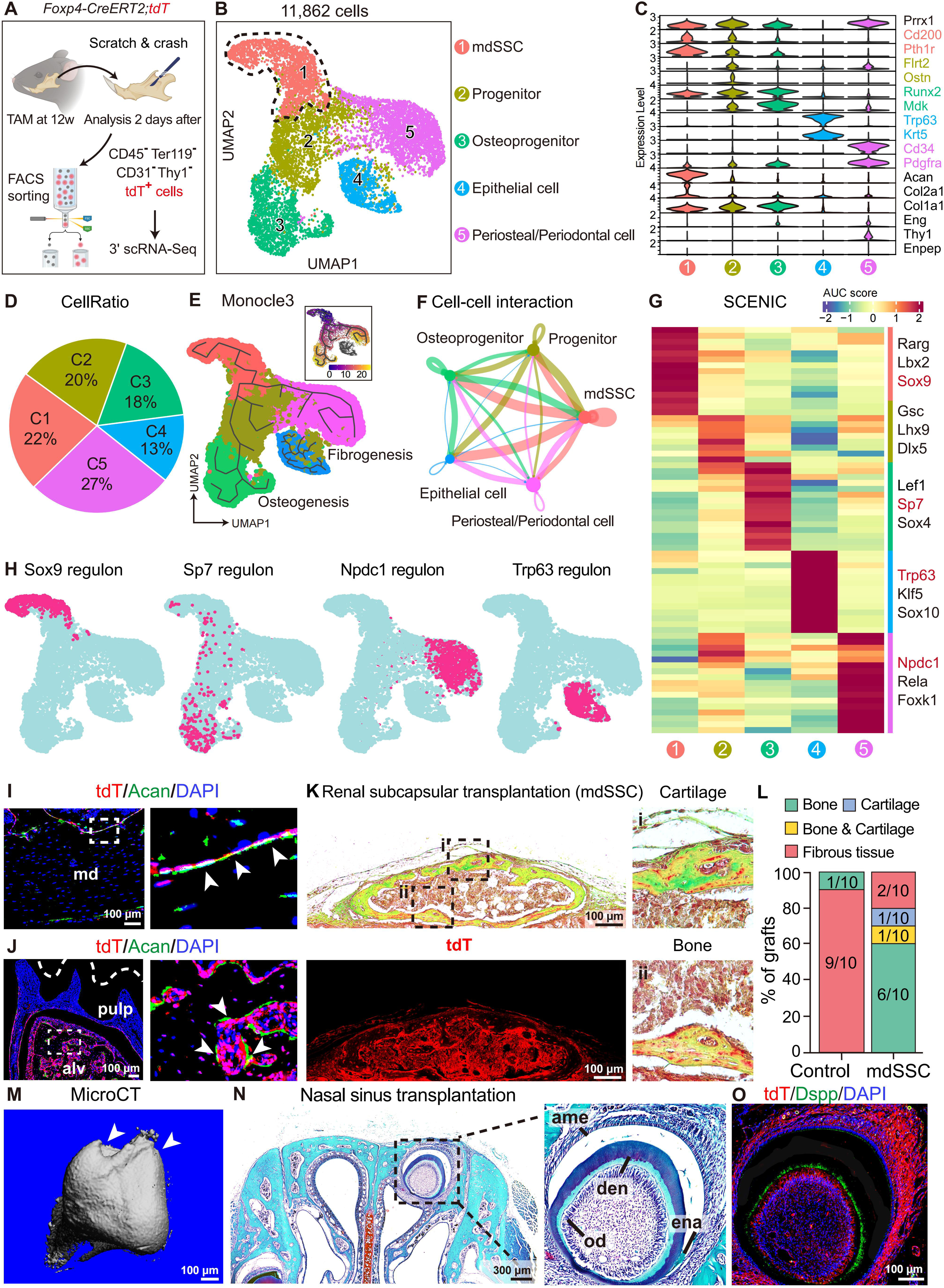
Identification and functional characterization of Foxp4^+^ mdSSCs in adult mouse mandible. (A) scRNA-seq workflow and experimental scheme. (B) UMAP visualization of 5 cell clusters in 12-week-old adult mandible. In total, 11,862 single cells from 6 mice were analyzed. Dotted lines indicated the mdSSC cluster. (C) Violin plots showing the expression of feature genes across the 5 cell clusters. Marker genes for annotating each cell cluster were color-coded on the right. (D) Pie chart showing the cell ratios of each cell cluster. (E) UMAP visualization of the osteogenesis and fibrogenesis trajectories simulated by Monocle3. Diffusion pseudotime was indicated in the upper right panel. (F) Circle plot showing cell-cell interactions among the 5 cell clusters. The interaction strength (indicated by line thickness) was calculated by CellChat. (G) Heatmap showing regulons enriched in each cell cluster. The AUC score (row scaling) was computed. Three representative regulons (out of top 10 regulons) for each cell cluster were listed on the right. (H) Binary activities of Sox9, Sp7, Npdc1 and Trp63 regulons were shown by UMAP plots. (I and J) IF images of Foxp4^+^ mdSSCs in adult mandibular (I) and alveolar (J) bone. Tamoxifen was administered for 5 consecutive days in 12-week-old *Foxp4-CreERT2; tdTomato* mice and analyzed 1 day after. The tdTomato (tdT), Acan and DAPI signals were shown. Dotted lines (J) indicated the border of the dental crown (n = 3 mice per group from 3 independent experiments). Arrowheads indicated the tdT^+^Acan^+^ mdSSCs. md: mandibular bone, alv: alveolar bone. (K) Movat’s pentachrome staining showing cartilage (i) and bone (ii) formation after renal subcapsular transplantation of adult Foxp4^+^ mdSSCs from 12-week-old *Foxp4-CreERT2; tdTomato* mice. The tdTomato (tdT) signal (lower left panel) demonstrated donor-derived cells. (L) Quantification of the renal subcapsular transplantation data (n=10 mice per group from 8 independent experiments). (M) Representative microCT image of the ectopic tooth formed by adult Foxp4^+^ mdSSCs 4 weeks after nasal sinus transplantation. Arrowheads indicated the tooth cups. (N and O) Safranin O/Fast Green (N) and IF (O) staining of the ectopic tooth formed by adult Foxp4^+^ mdSSCs 4 weeks after nasal sinus transplantation. Foxp4^+^ mdSSCs from 12-week-old *Foxp4-CreERT2; tdTomato* mice were transplanted.

Since Acan is more specifically expressed in the mdSSC subset, we performed IF staining of Acan to detect the localization of Foxp4^+^ mdSSCs in 12-week-old *Foxp4-CreERT2; tdTomato* mandible. Consistent with the fact that adult mandible does not contain cartilage tissues,^71^ tdTomato^+^ Acan^+^ cells mainly distributed in the endosteal region of the mandibular bone (Figure 4I). In 12-week-old molars, tdTomato^+^ Acan^+^ cells were also found in the endosteal region of the alveolar bones (Figure 4J).

To test the stem cell activity of Foxp4^+^ mdSSCs in adult mouse mandible, we administered tamoxifen in 12-week-old *Foxp4-CreERT2; tdTomato* mice for 5 consecutive days (Figure S4A), flow cytometrically sorted CD45^-^ Ter119^-^ CD31^-^ Thy1^-^ 6C3^-^ CD200^+^ CD105^-^ cells from molar-extracted mandibles 2 days later, and sub-divided them into tdTomato^+^ (mdSSCs) or tdTomato^-^ (control) fractions (Figure S4B). CFU-F assay showed significantly higher clonogenic activity of mdSSCs (Figure S4C). *In vitro* trilineage differentiation analyses showed significantly increased osteogenic, adipogenic and chondrogenic capacities of mdSSCs as compared to control cells (Figures S4D-S4F). To test the *in vivo* osteo-chondrogenic potential of 12-week-old mdSSCs, we performed renal subcapsular transplantation (Figure S4G) and found that mdSSCs gave rise to more ectopic bone and cartilage tissues as compared to control cells (Figures 4K, 4L and S4H). To test the odontogenic potential of 12-week-old mdSSCs, we performed nasal sinus transplantation (Figure S4G) and found that they could regenerate ectopic tooth with multi-cusp dental crown (Figure 4M), pulp, dentin and enamel structures (Figures 4N and 4O) after 4 weeks. Nasal transplantation of 12-week-old mdSSCs also generated ectopic bone, but not cartilage, while control cells mainly generated fibrous tissue as revealed by histological analyses (Figures S4I-S4L). To test the maturity and stability of the regenerated tooth, we performed microCT analysis 8 weeks after nasal sinus transplantation, and found evident dental root elongation with root apex, suggesting continued maturation of the ectopic tooth (Figure S4M). Therefore, we identified and functionally characterized Foxp4^+^ mdSSCs in adult mouse mandibles.

### Genetic ablation of Foxp4^+^ cells impairs mandible development and fracture repair

To test whether Foxp4^+^ cells are required during mandible development and repair, we crossed the *Foxp4-CreERT2; tdTomato* and *Rosa26-DTA* alleles to generate *Foxp4-CreERT2; tdTomato; DTA* mice (DTA mice hereafter), in which Foxp4^+^ cells could be conditionally ablated by diphtheria toxin. We first administered tamoxifen at E15.5 and analyzed mandibular bone and tooth development at E19.5 (Figure 5A). DTA mice showed significantly reduced mandible length and impaired molar tooth germ formation as compared to littermate control mice (*Foxp4-CreERT2; tdTomato*, Figures 5B-5D). In contrast, no significant maxillary defects were observed (Figures 5B-5D), consistent with the fact that Foxp4 is more abundantly expressed in the developing mandible as compared to maxilla (Figures S1I and S1J). IF staining showed efficient ablation of tdTomato^+^ cells, as well as significantly reduced number of Col2a1^+^ chondrocytes and Runx2^+^ osteoprogenitors in DTA as compared to littermate control mice (Figures 5E-5J).

**Figure 5.**
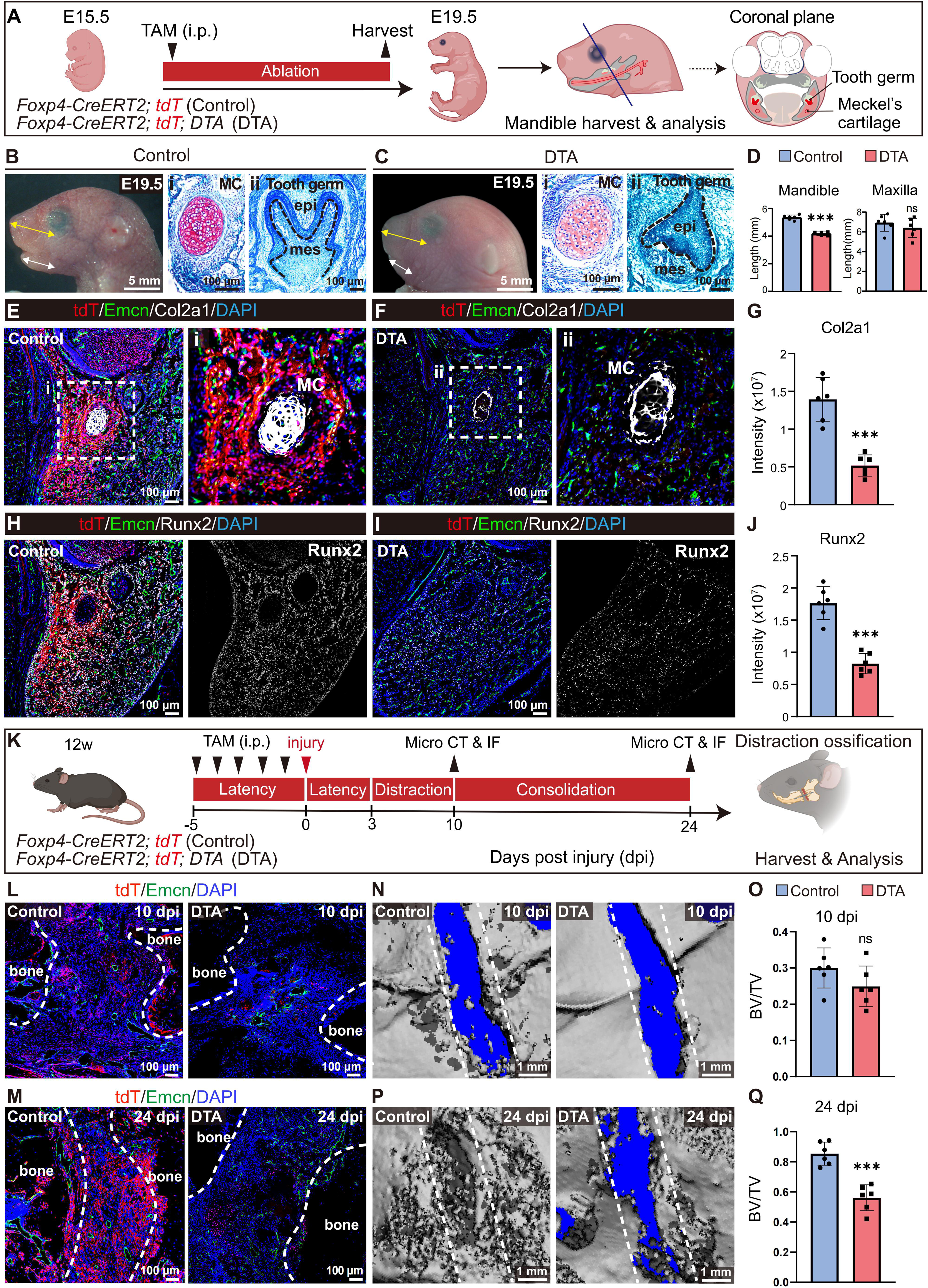
Ablation of Foxp4^+^ cells impairs mandibular development and fracture repair. (A) Schematic illustration of Foxp4^+^ cell ablation during mandible development. Tamoxifen was administered for 1 day at E15.5. *Foxp4-CreERT2; tdTomato* (Control) or *Foxp4-CreERT2; tdTomato; DTA* (DTA) mice were analyzed at E19.5. (B and C) Representative images showing mandibular and tooth germ defects associated with perinatal DTA mice. Lateral views of control or DTA skulls were shown, with double-headed arrows indicating the length of the developing mandible (white) and maxilla (yellow). Safranin O/Fast Green staining of MC (i) and tooth germ (ii) were also provided. Dotted lines (ii) indicated the border of odontogenic epithelium and tooth germ mesenchyme. mes: mesenchyme, epi: epithelium. (D) Quantification of the mandibular and maxillary length at E19.5 (n=6 mice per genotype from 3 independent experiments). (E-J) IF images of control and DTA mandibles at E19.5 with quantifications of Col2a1 (G) and Runx2 (J) expression. The tdTomato (tdT), Emcn, Col2a1, Runx2 and DAPI signals were shown. MC: Meckel’s cartilage. Quantification of the fluorescence signals was performed by Image J (n=6 mice per genotype from 3 independent experiments). (K) Schematic illustration of Foxp4^+^ cell ablation during mandibular fracture repair in adult mice. Tamoxifen was administered consecutively for 5 days before DO surgery at 12-week-old. After 3 days of latency period, *Foxp4-CreERT2; tdTomato* (Control) or *Foxp4-CreERT2; tdTomato; DTA* (DTA) mice were analyzed at 10 and 24 dpi. (L and M) IF images of control and DTA mandibular bone callus at 10 and 24 dpi. The tdTomato (tdT), Emcn and DAPI signals were shown. Dotted lines indicated the osteogenic front. (N-Q) MicroCT images of control and DTA mandibular bone callus at 10 (N) and 24 (P) dpi with quantifications (O and Q). BV/TV: bone volume ratio. Dotted lines indicated the fracture sites. Statistical significance was determined by unpaired two-tailed Student’s t test. All data represent mean ± SD (ns: not significant, ^∗∗∗^p < 0.001).

To test whether Foxp4^+^ cells are required for adult mandible repair after fracture, we administered tamoxifen in 12-week-old DTA or control mice before DO surgery (Figure 5K). IF staining demonstrated efficient ablation of tdTomato^+^ cells at 10 and 24 dpi (Figures 5L and 5M). MicroCT analysis of the bone callus showed significantly decreased bone volume ratio in DTA mice as compared to littermate controls at 24 dpi (Figures 5N-5Q). Taken together, we found that genetic ablation of Foxp4^+^ cells impairs both mandible development and fracture repair in adult mice.

### Primary cilia are indispensable for Foxp4^+^ mdSSCs to promote mandible development and fracture repair

To explore the mechanisms by which Foxp4^+^ mdSSCs are regulated during mandible development and regeneration, we analyzed the scRNA-seq datasets in both E12.5 and 12-week-old mice (Figures 2B and 4B). Gene ontology (GO) analysis revealed that genes related to “Response to mechanical stimulus” were enriched in mdSSCs at both time points, and that genes related to “Cilium assembly” was enriched in 12-week-old mdSSCs (Figures S5A and S5B). KEGG analysis showed enrichment of calcium, Wnt and Hedgehog signaling pathways in E12.5 and adult mdSSCs, all of which are closely related to primary cilia function^72^ (Figures S5C and S5D). Bulk RNA-seq in E12.5 mandible showed that Foxp4^+^ mdSSCs enriched genes related to “Cilium organization” and “Cilium assembly” as compared to control cells (Figures S5E and S5F), in which *Ift140* was the most significantly up-regulated intraflagellar transport (IFT) gene that facilitates retrograde transport within the primary cilia (Figure S5G).^73^

To test whether Ift140 regulates mdSSC function, we first performed siRNA-mediated knockdown of *Ift140 in vitro*, and found that *Ift140* silencing led to significant down-regulation of Wnt and Hedgehog signaling target genes (Figure S5H). CFU-F and trilineage differentiation analyses showed that *Ift140* silencing impairs both colony-forming and differentiation capacities of mdSSCs (Figures S5I-S5L). Next, we crossed the *Foxp4-CreERT2* and *Ift140^flox^* alleles to generate *Foxp4-CreER; Ift140^flox/flox^* mice (cKO mice), in which primary cilia formation and function are conditionally impaired.^74,75^ We first administered tamoxifen at E15.5 and analyzed their perinatal phenotypes at E19.5 (Figure 6A). Similar to DTA mice, cKO mice show mandibular, but not maxillary, shortening and delayed molar tooth germ development (Figures 6B-6D). IF staining showed significantly reduced number of primary cilia and Foxp4^+^ cells in cKO as compared to control mice (Figures 6E-6J). Col2a1^+^ chondrocytes and Runx2^+^ osteoprogenitors were also significantly reduced in cKO mice (Figures 6K-6P).

**Figure 6.**
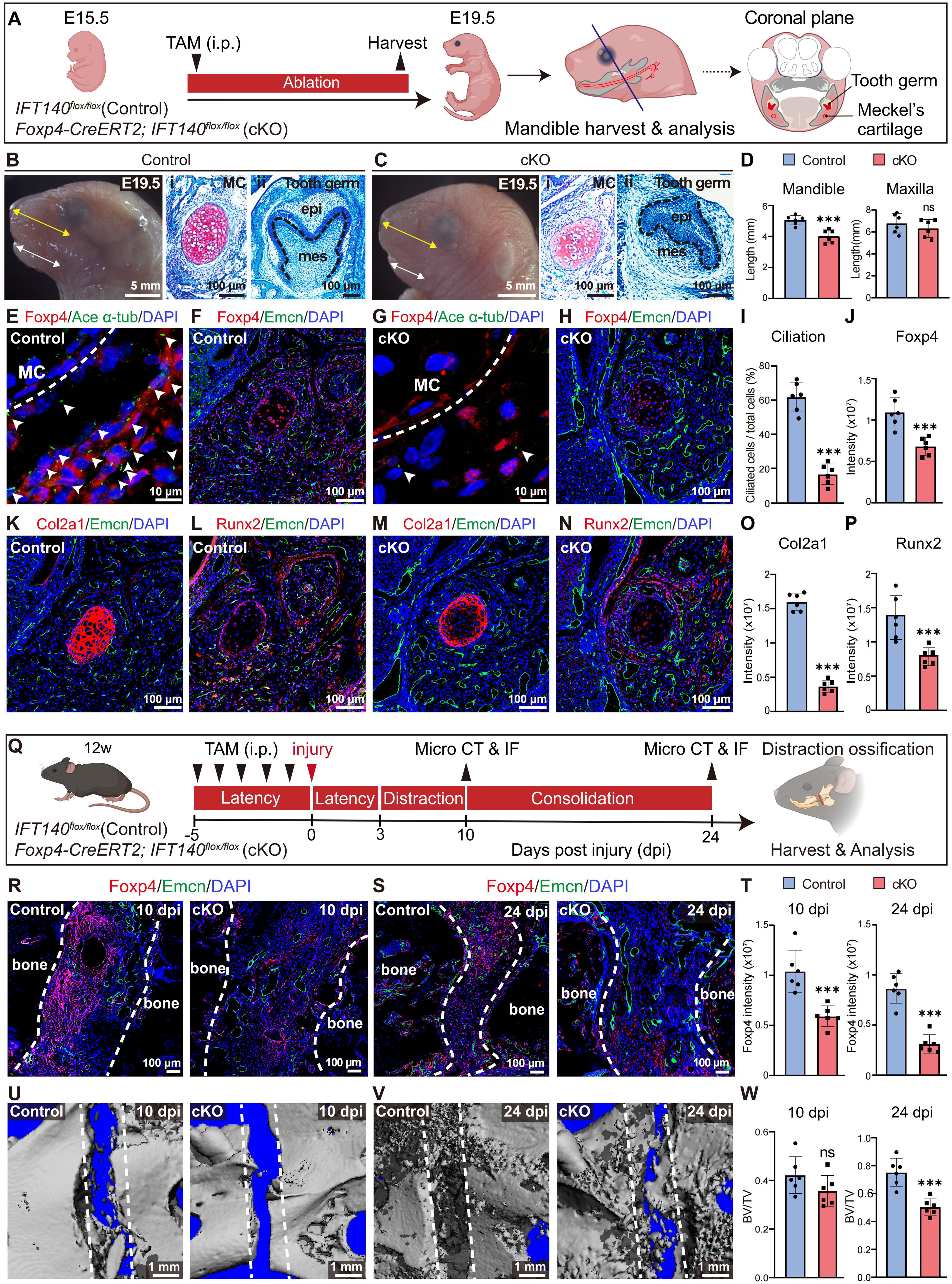
Ddeletion of *Ift140* from Foxp4^+^ cells impairs primary cilia formation, mandible development and fracture repair. (A) Schematic illustration of *Ift140* conditional deletion during mandible development. Tamoxifen was administered for 1 day at E15.5. *Ift140^fl/fl^* (Control) and *Foxp4-CreERT2; Ift140^fl/fl^* (cKO) mice were analyzed at E19.5. (B and C) Representative images showing mandibular and tooth germ defects associated with perinatal cKO mice. Lateral views of control or cKO skulls were shown, with double-headed arrows indicating the length of the developing mandible (white) and maxilla (yellow). Safranin O/Fast Green staining of MC (i) and tooth germ (ii) were also provided. Dotted lines (ii) indicated the border of odontogenic epithelium and tooth germ mesenchyme. mes: mesenchyme, epi: epithelium. (D) Quantification of mandibular and maxillary length at E19.5 (n=6 mice per genotype from 3 independent experiments). (E-J) IF images of control and cKO mandibles at E19.5 with quantifications of ciliated cells (I) and Foxp4 expression (J). The Foxp4, Ace α-tub (acetylated α-tubulin, primary cilia marker), Emcn and DAPI signals were shown. Arrowheads indicated primary cilia. Dotted lines indicated the border of Meckel’s cartilage (MC). Quantification of the fluorescence signals was performed by Image J (n=6 mice per genotype from 3 independent experiments). (K-P) IF images of control and cKO mandibles at E19.5 with quantifications of Col2a1 (O) and Runx2 (P) expression. The Col2a1, Runx2, Emcn and DAPI signals were shown. Quantification of the fluorescence signals was performed by Image J (n=6 mice per genotype from 3 independent experiments). (Q) Schematic illustration of *Ift140* conditional deletion during mandibular fracture repair in adult mice. Tamoxifen was administered consecutively for 5 days before DO surgery at 12-week-old. *Ift140^fl/fl^* (Control) and *Foxp4-CreERT2; Ift140^fl/fl^* (cKO) mice were analyzed at 10 and 24 dpi. (R-T) IF images of control and cKO mandibular bone callus at 10 (R) and 24 (S) dpi with quantification of Foxp4 expression (T). The Foxp4, Emcn and DAPI signals were shown. Dotted lines indicated the osteogenic front. bone: newly formed woven bone (n=6 mice per genotype from 3 independent experiments). (U-W) MicroCT images of control and cKO mandibular bone callus at 10 (U) and 24 (V) dpi with quantifications (W). BV/TV: bone volume ratio. Dotted lines indicated the fracture sites (n=6 mice per genotype from 3 independent experiments). Statistical significance was determined by unpaired two-tailed Student’s t test. All data represent mean ± SD (ns: not significant, ^∗∗∗^p < 0.001).

To test whether primary cilia regulate mandibular bone repair by Foxp4^+^ cells, we administered tamoxifen in 12-week-old cKO or control mice before DO surgery and analyzed their phenotypes at 10 or 24 dpi (Figure 6Q). Compared to littermate control mice, IF staining showed significantly reduced number of Foxp4^+^ cells in cKO mice at both 10 and 24 dpi (Figures 6R-6T). MicroCT analysis of the bone callus showed significantly decreased bone volume ratio in cKO mice as compared to littermate controls at 24 dpi (Figures 6U-6W). Importantly, transplantation of sphere-cultured^76^ mdSSCs into the fracture site significantly accelerated fracture repair in a Ift140-dependent manner (Figures S6A-S6I), further suggesting that primary cilia are indispensable for mdSSC activation and differentiation.

To test determine whether Foxp4 regulates primary cilia-associated signaling, we performed siRNA-mediated knockdown of *Foxp4* in cultured E12.5 mdSSCs and found that *Foxp4* silencing significantly down-regulates Hedgehog signaling target genes such as *Gli1/2* (Figure S6J). In contrast, Wnt signaling target genes such as *Dkk1* and *Axin2* were modestly upregulated (Figure S6J).

### Human FOXP4^+^ mdSSCs regenerate ectopic bone, cartilage and tooth after xenotransplantation

To test whether mdSSCs exist in adult human mandible, we re-analyzed our published scRNA-seq dataset of human mandibular bone fragments after mandibulectomy (Figures 7A and 7B).^77^ Interestingly, we found that cluster 4 highly expressed eSSPC marker *CADM1*,^44^ as well as hSSC markers *PDPN*, *CD164* and *CD73* (*NT5E*),^39^ suggesting a human mdSSC population (Figure 7C). Consistent with this hypothesis, gene regulatory network analysis showed that FOXP4, SOX9, GLI3 and TWIST1^78,79^ regulons were highly enriched in cluster 4 (Figures 7D and 7E). GO and KEGG analyses showed that genes related to “Response to mechanical stimulus”, “Cilium organization”, Wnt and Hedgehog signaling pathways were enriched in cluster 4 (Figures 7F and 7G). Therefore, we reasoned that mdSSCs could also exist in adult human mandibles.

**Figure 7.**
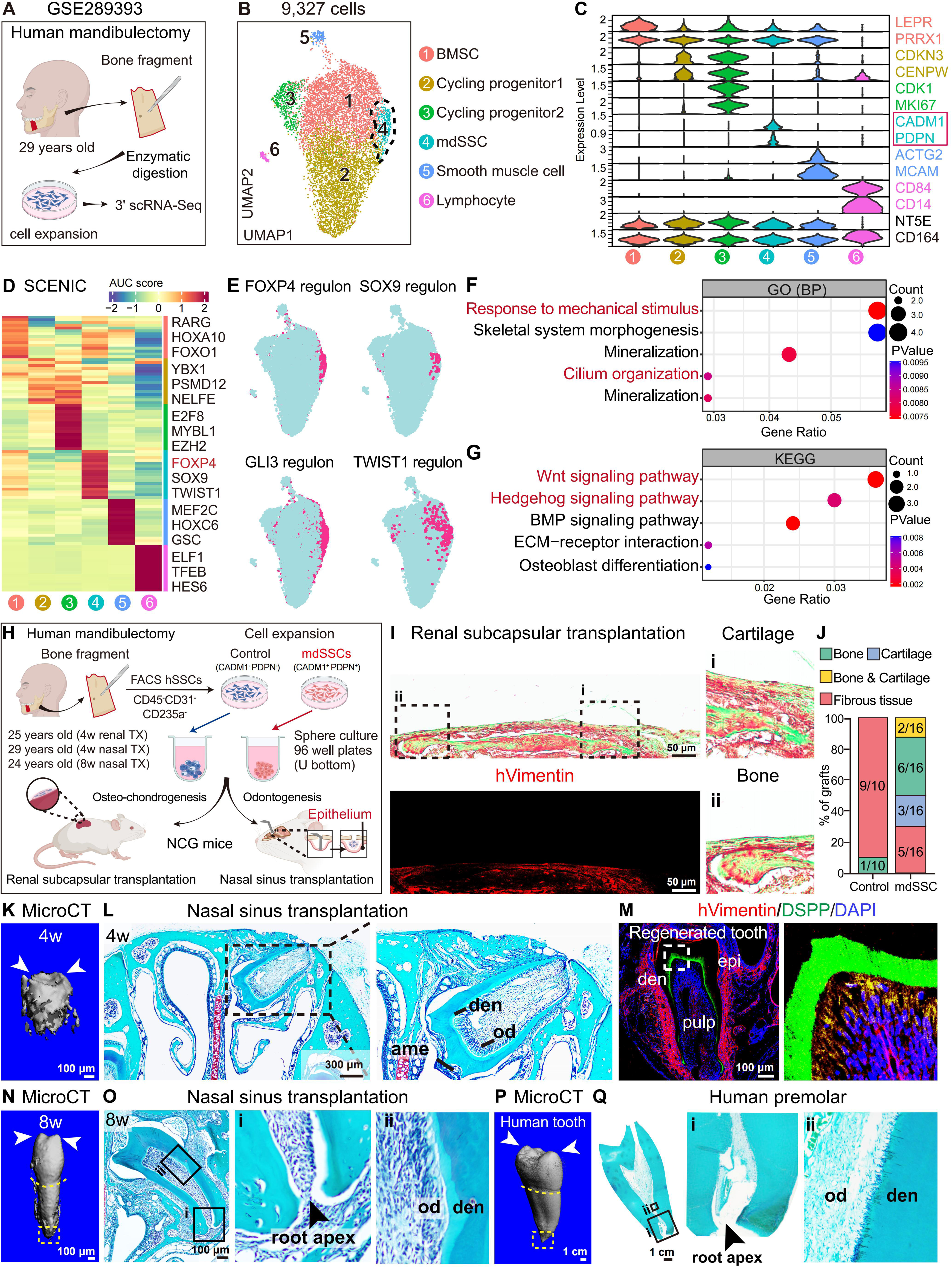
Identification and functional characterization of human FOXP4^+^ mdSSCs. (A) scRNA-seq workflow and experimental scheme. The mandibular bone fragment obtained from a 29-year-old patient was analyzed. (B) UMAP visualization of the 6 cell clusters in a previously published human mandible scRNA-seq data (GSE289393). In total, 9,327 cells were analyzed. Dotted lines indicated the mdSSC cluster. (C) Violin plots showing the expression of feature genes across the 6 cell clusters. Marker genes for annotating each cell cluster were color-coded on the right. (D) Heatmap showing the regulons enriched in each cell cluster. The AUC score (row scaling) was computed. Three representative regulons (out of top 10 regulons) for each cell cluster were listed on the right. (E) Binary activities of FOXP4, SOX9, GLI3 and TWIST1 regulons were shown by UMAP plots. (F and G) Dot plots showing GO and KEGG terms enriched in the mdSSC subset. BP: Biological process. (H) Transplantation workflow and experimental scheme. The mandibular bone fragments obtained from 24 to 29-year-old patients were analyzed. (I) Movat’s pentachrome staining showing cartilage (i) and bone (ii) formation after renal subcapsular transplantation of human mdSSCs. Lower left panel: IF images of human Vimentin (hVimentin) to demonstrate donor-derived cells. (J) Quantification of the renal subcapsular transplantation data (n=10 or 16 recipient mice per group from 5 independent experiments). (K) Representative microCT image of the ectopic tooth formed by human mdSSCs 4 weeks after nasal sinus transplantation into NCG mice. Arrowheads indicated the tooth cups. (L and M) Safranin O/Fast Green (L) and IF (M) staining of the ectopic tooth formed by human mdSSCs 4 weeks after nasal sinus transplantation into NCG mice. ame: ameloblast, den: dentin, od: odontoblast, epi: epithelium. Human Vimentin, DSPP and DAPI signals were shown in (M). (N) Representative microCT image of the ectopic tooth formed by human mdSSCs 8 weeks after nasal sinus transplantation into NCG mice. Arrowheads indicated the tooth cups. Dotted line indicated the boundary between dental crown and root. Dotted box indicated the root apex. (O) Safranin O/Fast Green staining of the ectopic tooth formed by human mdSSCs 8 weeks after nasal sinus transplantation into NCG mice. den: dentin, od: odontoblast. Arrowhead indicated the root apex. (P) Representative microCT image of a 14-year-old human premolar. Arrowheads indicated the tooth cups. Dotted lines indicated boundary between dental crown and root. Dotted box indicated the root apex. (Q) Safranin O/Fast Green staining of the 14-year-old human premolar. Arrowhead indicated the root apex. den: dentin, od: odontoblast.

To test this, we crushed and enzymatically digested adult human bone fragments obtained by mandibulectomy (Figure S7A), and sorted CD45^-^ CD31^-^ CD235a^-^ CADM1^+^ PDPN^+^ mdSSCs together with CD45^-^ CD31^-^ CD235a^-^ CADM1^-^ PDPN^-^ control cells by flow cytometry (Figure S7B). qPCR analysis confirmed that *FOXP4* expression is highly enriched in mdSSCs (Figure S7C). Similar to mouse mdSSCs, adult human mdSSCs showed significantly higher CFU-F forming activity and trilineage differentiation capacities (Figures S7D-S7G). Next, we performed xenotransplantation into immunodeficient mice (NCG) after 2D expansion and 3D cell sphere formation (Figure 7H). Compared to control cells that mainly generated fibrous tissue, adult human mdSSCs gave rise to bone and cartilage tissue upon renal subcapsular transplantation (Figures 7I and 7J). To test whether adult human mdSSCs retain odontogenic capacity, we performed nasal sinus transplantation and found that mdSSCs, but not control cells, generated ectopic human tooth after 4 weeks (Figures 7K and 7L), the origin of which was confirmed by human Vimentin staining (Figures 7M and S7H). We also detected ectopic bone formation after nasal sinus transplantation (Figures S7I-S7K).

To further test the maturity and stability of the regenerated tooth, we performed microCT and histological analyses 8 weeks after nasal sinus transplantation. Notably, the regenerated tooth exhibited a distinct morphology similar to a 14-year-old human premolar (Figures 7N-7Q). The root of the regenerated tooth was significantly elongated as compared to 4 weeks after transplantation, with a closed root apex (Figures 7N and 7O). A clear boundary between the root and multilZIcusp crown was also evident (Figure 7N). Collectively, these data demonstrated continued maturation of the regenerated tooth over time, and its morphological resemblance to human premolar.

To determine whether *FOXP4* is required for maintaining the clonogenic and multilineage differentiation capacities, we knocked it down in human mdSSCs and performed functional characterizations both *in vitro* and *in vivo*. *FOXP4* silencing significantly reduced the number of CFU-F colonies formed by human mdSSCs, compromised their *in vitro* differentiation (Figures S7L-S7P), and abolished *in vivo* tooth regeneration after nasal sinus transplantation (Figure S7Q). Taken together, we showed that mdSSCs are evolutionarily conserved in adult human mandible, which can regenerate ectopic bone, cartilage and tooth after xenotransplantation.

## Discussion

In this study, we identified Foxp4^+^ mdSSCs in both developing and adult mouse mandibles. In fetal mice, neural crest-derived Foxp4^+^ mdSSCs not only specify MC formation and promote mandibular osteogenesis, but also give rise to tooth germ mesenchyme to initiate molar development. In early postnatal and adult mice, Foxp4^+^ mdSSCs orchestrate osteogenesis and odontogenesis, and promote fracture repair during DO. Mechanistically, we found that primary cilia are indispensable for Foxp4^+^ mdSSCs during mandible development and regeneration. More importantly, we showed that FOXP4^+^ mdSSCs are evolutionarily conserved in adult human mandible, and that xenotransplantation of FOXP4^+^ mdSSCs into immunodeficient mice successfully regenerate human bone, cartilage and structurally intact tooth *in vivo.* Together, we discovered a mandible-derived SSC population that could be harnessed to accelerate fracture healing in mandibular bones, and to achieve whole tooth regeneration by stem cell therapy.

The appearance of mandible is an evolutionary milestone, since it promotes chewing efficiency and broadens the dietary repertoire of the modern vertebrates.^80^ Whereas previous studies have identified region-specific SSCs in the bone marrow, periosteum, growth plate, cranial suture and vertebral body,^34,38,40,42,81,82^ whether the mandible contains functionally distinct SSCs remains unknown. This uncertainty is primarily due to the unique feature of mandible development, which involves complex interplay between bone, cartilage and tooth. Here, we explored the stem cell basis of mandible development, and found that Foxp4^+^ mdSSCs orchestrate MC, bone and tooth germ formation. Genetic lineage tracing and scRNA-seq analyses revealed that Foxp4^+^ mdSSCs emerged around E12.5 to initiate MC and tooth germ development. However, the biochemical and mechanical cues that specify their distinct cell fates remain to be explored. Previous study showed that the anterior part of the mandible is formed by endochondral ossification, while the posterior part is formed by an undefined mechanism.^83,84^ In this study, we found that Foxp4^+^ mdSSCs first generate the MC template, which then localize in the perichondrial regions to promote osteogenesis and MC degradation (without cell invasion and bone marrow cavity formation). This special way of ossification is reminiscent of perichondral ossification,^83,85–87^ which is distinguishable from classical intramembranous or endochondral ossification.^88,89^ Notably, the posterior part of MC also generates malleus and incus,^90,91^ suggesting that Foxp4^+^ mdSSCs might also contribute to middle ear formation.

Another important finding of this study is that Foxp4^+^ mdSSCs also exist in postnatal mandibles. Although mature mandibles are devoid of cartilage tissue, adult mdSSCs highly express cartilage matrix proteins such as Acan and Col2a1, and the master chondrogenic transcription factor Sox9.^92^ This is in line with previous findings that postnatal SSCs highly express cartilage signature genes,^93–95^ and suggests that adult mdSSCs could be the remnants of fetal mdSSCs that retain osteo-chondrogenic capacity. Importantly, we found that adult mandible fracture induced by DO reactivates osteo-chondrogenic differentiation by Foxp4^+^ cells to promote fracture repair, while genetic ablation of Foxp4^+^ cells or their primary cilia severely delayed fracture healing. Furthermore, transplantation of cultured Foxp4^+^ mdSSCs promotes DO repair in a primary cilia-dependent manner. Since the primary cilia regulate diverse cellular functions^74^ as to mechanosensing, signal transduction, chemotaxis and cell polarity, further studies are needed to elucidate the detailed mechanisms by which primary cilia finetunes quiescence and activation of Foxp4^+^ mdSSCs. In the future, pharmacological activation of endogenous Foxp4^+^ mdSSCs (eg. by stimulating signaling pathways transmitted by primary cilia) could be an alternative approach to accelerate DO-induced mandible repair in clinical settings.

*In vivo* regeneration of intact and functional tooth has long been a huge challenge, since tooth formation requires complex interplay between odontogenic epithelium and neural crest-derived mesenchyme cells.^26,55^ Previous tooth regeneration strategies combine dental epithelial cells with mesenchymal cells (either intact dental mesenchyme or enzymatically dissociated cells) before transplantation into the subcutaneous or renal subcapsular depots, in which the transplanted mesenchymal cells are rather heterogeneous.^96–98^ Our study differs fundamentally from these approaches in that we transplanted highly purified, phenotypically defined mdSSCs without adding dental epithelial cells. Inspired by the similar properties shared between oral and nasal sinus epithelium and the clinical observation that supernumerary tooth can be found in the nasal cavity under certain pathological conditions,^57–59^ we pioneered the nasal sinus transplantation system that enables reprogramming of nasal sinus epithelium into odontogenic epithelium by mdSSCs. This system allows accurate assessment of the odontogenic potential of mdSSCs at different developmental stages. Remarkably, either embryonic (E12.5 and E15.5-E16.5) or adult (mouse and human) mdSSCs could regenerate ectopic tooth upon nasal sinus transplantation, suggesting that Foxp4^+^ mdSSCs persists in the mandible from early embryonic stages through adulthood. As far as we are concerned, loss of odontogenic epithelium over time underlies diminishing tooth formation capacity during mandible development and postnatal growth. In the future, it would be intriguing to test whether SSCs isolated from other parts of the adult skeleton could regenerate tooth-like structure upon nasal sinus transplantation.

Another exciting direction would be *in situ* regeneration of native whole tooth. This could be achieved by transplanting cultured Foxp4^+^ mdSSCs, or ectopic tooth germ formed by Foxp4^+^ mdSSCs in the nasal sinus, into the alveolar socket. Alternatively, a tooth germ organoid could be engineered *in vitro* by co-culturing oral/nasal epithelial cells and Foxp4^+^ mdSSCs before *in situ* transplantation. Notably, we found that human FOXP4^+^ mdSSCs obtained from human mandibular bone fragments can regenerate intact human whole tooth, which contain dental crown with cusps, dental root with apex, dental pulp and periodontal tissues, after xenotransplantation into the nasal sinus of immunodeficient mice. In clinical settings, FOXP4^+^ mdSSCs could be isolated from mandibular bones after wisdom tooth removal, followed by autologous transplantation to regenerate the loss tooth in other positions. Whereas FOXP4^+^ mdSSCs is a promising stem cell source for clinical application, the morphology of the regenerated tooth is governed by both biochemical and biomechanical microenvironments. Therefore, advanced bioengineered scaffolds as well as proper mechanical stimulations are indispensable to translate the discovery of this study into regenerative therapies in the near future.

### Limitations of the study

Although we successfully demonstrated that Foxp4^+^ mdSSCs exhibit odontogenic activity following ectopic transplantation into the nasal sinus, *in situ* (orthotopic) tooth regeneration within the native mandibular microenvironment (alveolar bone) has not been achieved, which is a critical next step for clinical translation. Furthermore, the efficiency of tooth formation as well as the morphology of the regenerated tooth remain to be optimized by future studies. Whereas we showed that primary cilia tightly regulate mdSSC function, other mechanisms might also play important roles in governing cell fate determination of Foxp4^+^ mdSSCs during mandibular/tooth development and regeneration, which awaits elucidation by future studies in order to translate the mdSSC-based cell therapy into clinical applications.

## ACKNOWLEDGMENTS

This work was supported by grants from the National Natural Science Foundation of China (32425027, 32330050, 82425014, U23A20444, 82270963, 82361148131, 82302706), National Key R&D Program of China (2022YFA1103200, 2021YFA1100900), Shanghai Municipal Science and Technology Commission (23XD1423900), Peak Disciplines (Type IV) of Institutions of Higher Learning in Shanghai, Shanghai Pilot Program for Basic Research, Shanghai Municipal Education Commission Scientific Research and Innovation Program (2023ZKZD29), and Fundamental and Interdisciplinary Disciplines Breakthrough Plan of the Ministry of Education of China (JYB2025XDXM508). R.Y. is a SANS exploration scholar.

## AUTHOR CONTRIBUTIONS

L.Z. and X.L. performed most of the experiments. D.C. performed bioinformatic data analysis. H.Y. helped with nasal sinus transplantation, flow cytometry and cell culture experiments. G.L. and P.N. collected human mandibular samples. J.W. and Q.Z. designed and characterized *Foxp4-CreERT2* mice. X.Z. and XY.X. helped with genotyping, *in vitro* differentiation, and renal subcapsular transplantation. XQ.X. and X.G. helped with mouse mandibular DO surgery. C.C. and D.T. helped with nasal sinus transplantation. X.W. and M.Y. critically read the manuscript and gave helpful suggestions. Y.S. provided *Ift140^flox^*mice, human mandibular samples, and supervised the study. R.Y. invented nasal sinus transplantation, designed and interpreted all experiments, and wrote the manuscript.

## STAR METHODS RESOURCE AVAILABILITY

### Lead contact

Further information and requests for resources and reagents should be directed to and will be fulfilled by the lead contact, Rui Yue.

### Materials availability

All biological materials used in this study are available from the lead contact upon request or from commercial sources.

### Data and code availability

- scRNA-seq data have been deposited to Gene Expression Omnibus (GEO) database (GSE27786). Bulk RNA-seq data have also been deposited to GEO database (GSE324096).
- This paper does not report original code. We use referenced sources of code that can be found in the vignettes of the cited packages as listed in the method.
- Any additional information required to reanalyze the data reported in this paper is available from the lead contact upon request.

## EXPERIMENTAL MODEL AND SUBJECT DETAILS

### Mouse strains

All mice were maintained on C57BL/6J background, including *Foxp4-CreERT2, Ift140^flox^,*^99^ *Rosa26-tdTomato*,^100^ *Rosa26-DTA*,^101^ NCG (GemPharmatech, T001475) and wild-type mice. The *Foxp4-CreERT2* knock-in mice were generated by CRISPR/Cas9 technology. A *CreERT2-IRES-EGFP-WPRE-PolyA* cassette was inserted right after the translational start site (ATG) of the endogenous *Foxp4* gene locus (C57BL/6J background, Shanghai Biomodel Organism). All procedures were approved by the Tongji University Animal Care and Use Committee. Animals were housed in the SPF facility of the Animal Resource Center at Tongji University, with a 12-hour light/12-hour dark photoperiod. Animals were randomly allocated to different experimental groups. The specific number and age of animals in each experiment was indicated in the corresponding figure legends.

### Human samples

This study was carried out in compliance with the Helsinki Declaration. All human samples were collected and processed in accordance with the official ethical guidelines and protocols approved by the Ethics Committee of the Shanghai Tongji Stomatological Hospital. Human mandibular bone fragments were collected from patients with craniofacial abnormalities during mandibulectomy.

## METHOD DETAILS

### Tamoxifen administration

Tamoxifen (MCE, HY-13757A) was dissolved in corn oil (20 mg/ml) and was administered (100 mg/kg body weight) by intraperitoneal injection at indicated time points. To specifically label, ablate or conditionally delete specific genes from Foxp4-expressing cells, *Foxp4-CreERT2; tdTomato,* DTA, cKO or control mice were administered with different dosages of tamoxifen as indicated in the corresponding figure legends.

### Genotyping

Mice tail tips were lysed using DirectPCR Lysis Reagent (Viagen Biotech, 102-T) according to the manufacturer’s instructions. PCRs were performed to identify mice with Cre recombinase and/or loxP sites. Primer sequences for genotyping can be found in Table S1. The size of the band obtained from mouse genotyping is: *Ift140^flox^*: wild type = 319 bp, heterozygote = 375 bp and 319 bp, homozygote = 375 bp; *Foxp4-CreERT2*: wild type = 512 bp, heterozygote = 528 bp and 512 bp, homozygote = 528 bp; *Rosa26-DTA:* wild type = 415 bp, heterozygote = 302 bp and 415 bp, homozygote = 302 bp; *Rosa26-tdTomato:* wild type = 297 bp, heterozygote = 196 bp and 297 bp, homozygote = 196 bp.

### Renal subcapsular transplantation

Renal subcapsular transplantation was performed as previously described.^102^ Eight to twelve-week-old wild-type mice (for mouse cells) or NCG immune-deficient mice (for human cells) were used as recipients. Briefly, mice were anaesthetized and shaved on the left flank and abdomen before sterilization at the surgical site. Ketoprofen was injected intraperitoneally both before and after the surgery (every 12 hours for up to 72 hours). The kidney was externalized through a 1 cm incision and a 2 mm pocket was made in the renal capsule. Three hundred thousand 2D-cultured mouse or human cells were pelletized in round bottom ultra-low attachment microplates for 2 days, and transplanted underneath the renal capsule with a Matrigel plug. The incision was sealed using a cauterizer before replacing the kidney back to the body cavity. Animals were euthanized by CO_2_ after 4 weeks of transplantation. Kidneys were isolated and fixed with 4% PFA for 24 hours at 4°C, followed by dehydration, paraffin embedding, sectioning and Movat’s pentachrome staining (ScyTek, MPS-1) to demonstrate bone and cartilage differentiation. Immunostaining was also performed on adjacent sections.

### Nasal sinus transplantation

Eight to twelve-week-old wild-type mice (for mouse cells) or NCG immunodeficient mice (for human cells) were used as recipients. Mice were anaesthetized and the nasal skin was shaved and scrubbed with 75% ethanol. Ketoprofen was injected intraperitoneally both before and after the surgery (every 12 hours for up to 72 hours). An incision was made by #11 blade. A 0.8 mm-diameter drill was used to make a hole on the left nasal bone without breaking the nasal epithelium. Three hundred thousand 2D-cultured mouse or human cells were pelletized in round bottom ultra-low attachment microplates for 2 days, and then transplanted on top of the nasal sinus epithelium with a Matrigel plug (Corning, 356238). The incision was closed using 6–0 sutures. Animals were euthanized by CO_2_ at 4 or 8 weeks after transplantation. The facial bones containing nasal sinus were isolated and fixed with 4% PFA for 24 hours at 4°C, followed by dehydration, paraffin embedding, sectioning and Safranin O/Fast Green to demonstrate tooth, bone and cartilage differentiation (Solarbio, G1371). IF staining was also performed on adjacent sections.

### Distraction ossification model

The mandibular DO surgery was performed as previously described.^7^ Mandibular distraction devices were manufactured by computer-aided design and 3D printing at 20 µm resolution (Unigraphics NX10.0, Bosheng Mould). Twelve-week-old mice were used for all bone injury models, which were anesthetized using Avertin. Ketoprofen was injected intraperitoneally both before and after the surgery (every 12 hours for up to 72 hours). The mandible was exposed by making a 1 cm incision on the skin and separating the masseter muscle. Two 0.6 mm holes were drilled (6 mm apart with the cutting site in the middle), and the mandibular bone was cut posterior to the third molar using a fissure bur (0.8 mm). Distraction plates were secured with insertion of tight-fit 0.8 mm screws. The muscle and skin were then closed. The DO protocol started with a 3-day latency period, followed by distraction for 7 days at a rate of 0.2 mm/12 h, and up to 35 days of consolidation.

For orthotopic transplantation, 300,000 2D-cultured E12.5 mdSSCs were pelletized in round bottom ultra-low attachment microplates for 2 days to form 3D cell spheres, and then transplanted into the injury site with a Matrigel plug (Corning, 356238) every day for up to 3 days after DO surgery.

### MicroCT analysis

After carefully removing the DO devices, mandibles were fixed in 4% PFA for 24 hours at 4°C, and stored in PBS. The mandibles were analyzed by microCT 50 (Scanco Medical) at an isotropic voxel size of 10 μm, with peak tube voltage of 70 kV and current of 0.114 mA. A three-dimensional Gaussian filter (s=0.8) with a limited, finite filter support of one was used to suppress noise in the images, and a threshold of 220–1000 was used to segment mineralized bone from the air and soft tissues. For DO analysis, trabecular parameters were measured by analyzing 300 slices of the bone callus (with a threshold of <280 to exclude normal mandibular bones) at the injury site.

### Histological analysis

For histological analysis, mandibles and facial bones containing the nasal sinus were decalcified in 10% EDTA (pH 7.4) at 4°C for 2-4 weeks. After dehydration through a graded ethanol series, specimens were embedded in paraffin and cut into 6 μm sections. For morphological evaluation, the sections were stained with H&E (Sangon Biotech) or Safranin O/Fast Green Kit (Solarbio) according to the manufacturer’s instructions.

### IF staining and confocal imaging

For IF staining, mandibles and facial bones containing the nasal sinus were embedded in OCT and sectioned at 10-μm thickness after decalcification. Antigen retrieval was performed using hyaluronidase, sections were then blocked in 5% horse serum in PBST (0.1% Triton X-100 in PBS) for 30 min, and incubated overnight at 4 °C with primary antibodies, including anti-Col2a1 (1:200, Cat# BA0533, Boster Biological), anti-Acan (1:200, Cat# BA2967-1, Boster Biological), anti-Emcn (1:200, Cat# sc-65495, Santa Cruz), anti-Dspp (1:200, Cat# abs127711, Absin Bioscience), anti-Osteorix (Sp7) (1:500, Cat# ab209484, Abcam), anti-Runx2 (1:500, Cat# ab192256, Abcam), anti-Foxp4 (1:500, Cat# ABE74, Sigma), anti-ace-α-tubulin (1:500, Cat# T6793, Sigma), anti-human VIMENTIN (1:200, Cat# ab8069, Abcam), anti-Foxp1 (1:200, Cat# ab16645, Abcam), and anti-Foxp2 (1:200, Cat# ab16046, Abcam), anti-Ctsk (1:200, Cat# sc-48353, Santa Cruz), and anti-Postn (1:200, Cat# A01378, Boster Biological). After washing with PBS for 3 times (5 min each), sections were incubated with secondary antibodies, including: Donkey anti-rabbit Alexa Fluor 488 (1:500, Cat# A-21206, Invitrogen), Donkey anti-rabbit Alexa Fluor 594 (1:500, Cat# A-32754, Invitrogen), Donkey anti-rabbit Alexa Fluor 647 (1:500, Cat# A-31573, Invitrogen), Donkey anti-mouse Alexa Fluor 594 (1:500, Cat# A-21202, Invitrogen) and Donkey anti-rat Alexa Fluor 488 (1:500, Cat# A-48269, Invitrogen). After washing with PBS for 3 times (5 min each), sections were stained with DAPI (1:1000, Sigma) and mounted using Anti-Fade Aqueous Mounting Medium (Southern Biotech). All IF images were acquired using Nikon Ti2-E/A1R confocal microscope. Imaris (Oxford instruments) and Image J (1.51K) were used for quantitative analyses.

### Tissue clearing and lightsheet microscopy

Tissue clearing was performed as described previously.^103^ Briefly, intact mandible of E19.5 mouse was isolated, and attached soft tissues were removed using forceps. The tissue was immersed in 4% PFA for 48 h. Decalcification was performed with 10% EDTA solution for 24 h. Decolorization was performed using 25% Quadrol (Sigma-Aldrich 122262) solution for 1 day. Delipidation was performed using gradient tert-butanol (tB) (Sigma Aldrich 360538) solution (30%, 50% and 70%) for 1 day per concentration. Dehydration was performed using tB-PEG (PEGMMA500) (Sigma-Aldrich 409529) solution for 2 days with daily medium change. The tissue was finally perfused with the benzyl benzoate (BB) (Sigma Aldrich B6630)-PEG clearing solution until the tissue turned transparent, which took at least 24 h. Samples were maintained in clearing solution for subsequent imaging. Fluorescence images were acquired using ZEISS lightsheet microscope (Lattice Lightsheet 7). Imaris microscopy image analysis software (Oxford instruments) was used for quantitative analysis.

### Isolation and culture of primary cells

For E12.5 mouse mandibles, tissues were dissected under the microscope and minced using microsurgical scissors before enzymatic digestion. For adult mouse mandibles, the facial skin and muscle were carefully removed by microsurgical scissors and forceps. After extracting the incisors and molars, the periosteum was scratched by surgical blade and digested with the mandibular bone fragments (crushed into 1 mm^3^ pieces). For human mandibular bone fragments, samples were harvested within an hour after surgery and washed twice with PBS containing 1% penicillin/streptomycin (P/S). The periosteum was scratched by surgical blade and digested with the mandibular bone fragments (crushed into 1 mm^3^ pieces). The enzymatic digestion solution contains: 3 mg/ml type I collagenase (Worthington, LS004197), 4 mg/ml Dispase II (Sigma, 04942078001) and 1 U/ml DNase I (Sigma, D4527) in HBSS with Ca^2+^ and Mg^2+^. After three rounds of enzymatic digestions in 37°C water bath (15 min per round), cells were resuspended in staining buffer (Ca^2+^ and Mg^2+^ free HBSS with 2% FBS and 1% P/S) with 2 mM EDTA to stop the digestion. Dissociated cells were then filtered, centrifuged, and flow cytometrically sorted. Cells were cultured with α-MEM (Gibco, C12571500BT), 20% FBS (Gibco 10270-106, Lot: 42F6480K), 10 µM ROCK inhibitor (MedChemExpress, HY-15720), and 1% P/S (Gibco, 15-140-122). Culture medium was changed on day 2 and every 3-4 days thereafter. When cell confluency reached 80%, cells were dissociated by TrypLE Express without phenol red (Thermo) and passaged. For CFU-F culture, 1000 flow cytometrically sorted cells were seeded in 6-well plate with the culture medium. On day 8, cells were fixed and stained with crystal violet to quantify the colony number, and dissociated by TrypLE Express to calculate the colony size. For 3D sphere culture, 3_×_10^5^ passaged cells were plated in a 96-well round bottom ultra-low attachment microplate (Corning, 7007) with culture medium for 2 days. Cultures were maintained at 37_°_C with 5% O_2_ and 5% CO_2_ to create a low oxygen environment that promotes cell survival and proliferation.

### Flow cytometry

Dissociated mouse cells were stained on ice in staining buffer (Ca2^+^ and Mg2^+^ free HBSS with 2% FBS) with the following antibodies: APC-CD31 (1:200, BioLegend, clone#: 390), APC-CD45 (1:200, BioLegend, clone#: 30F-11), APC-Ter119 (1:200, BioLegend, clone#: TER-119), APC-Thy1.2 (1:200, BioLegend, clone#: 53-2.1), PE/Cy5-CD105 (1:200, BioLegend, clone#: MJ7/18), and PE/Cy7-CD200 (1:200, BioLegend, clone#: OX-90). Dissociated human cells were stained on ice in staining buffer with the following antibodies: APC-CD31 (1:200, BioLegend, clone#: WM59), APC-CD45 (1:200, BioLegend, clone#: HI30), PerCP-eFluor710-PDPN (1:200, BioLegend, clone#: NZ-1.3), Biotin-CADM1 (1:200, MBL, clone#: 3E1), PE/Cyanine7-Streptavidin (1:800, BioLegend, Cat# 405206). DAPI were used for viable cell gating. Flow cytometry analyses were performed using BD FACS Aria II and BD FACS Diva. Flow cytometry sorting was performed on BD FACSAria II and FUTech SE420 cell sorter. Acquired raw data were further analyzed on FlowJo version 10.9.0.

### *In vitro* differentiation

Primary cells were cultured for 7 days and passaged into 24-well plates at a cell density of 1×10^4^/cm^2^. After confluence, cell medium was changed to adipogenic or osteogenic medium. The adipogenic medium contains DMEM with low glucose (Gibco), 10% FBS (Gibco, 10270-106, Lot: 42F6480K), 1% P/S (Gibco, 15-140-122), 0.5 μM isobutylmethylxanthine (Sigma, I7018), 60 μM indomethacin (Sigma, 17378), 5 μg/ml insulin (Sigma, I0305000), and 1 μM dexamethasone (Sigma, D2915). Adipogenic medium was changed every 3 days. Oil red O (Sigma, O0625) staining was performed on day 7. The osteogenic medium contains α-MEM (Gibco), 10% FBS (Gibco, 10270-106, Lot: 42F6480K), 1 mM sodium pyruvate, 1% P/S, 50 mg/ml L-ascorbic acid, 10 mM β-glycerophosphate, and 100 nM dexamethasone. Osteogenic medium was changed every 3 days. Alizarin red (Sigma, A5533) staining was performed on day 14. For chondrogenic differentiation, 2.5×10^5^ cells were centrifuged in 15-ml polypropylene tubes to form cell pellets. The chondrogenic medium contains 10 ng/ml recombinant TGF-b3 (PeproTech, 100-36E), 100 nM dexamethasone, 50 μg/ml ascorbic acid 2-phosphate (Sigma, A8960), 1 mM sodium pyruvate, 40 μg/ml proline (Sigma, P0380), 1% P/S, and 1× ITS cell culture supplement (Cyagen, 6.25 μg/ml bovine insulin, 6.25 μg /ml transferrin, 6.25 μg /ml selenous acid, 5.33 μg/ml linoleic acid), and 1.25 mg/ml BSA in high glucose DMEM. The medium was changed every 3 days. Chondrogenic differentiation efficiency was determined by cryosection of the cell pellets and toluidine blue staining on day 21.

### RNA extraction and qPCR

Cells were lysed in RNAiso Plus (Takara) for total RNA extraction. RNA was reverse transcribed into cDNA using HiScript III RT SuperMix (Vazyme, R323-01). qPCR was performed using ChamQ Universal SYBR qPCR Master Mix (Vazyme, Q711-02) on a CFX96 real-time system (BioRad). Relative mRNA expression was calculated by normalizing to mouse β*-actin* or human *GAPDH*. For qPCR data analysis, the 2^−ΔΔCt^ method was used to calculate the relative fold-change of gene expression. The qPCR primer sequences are listed in Table S1.

### scRNA-seq library construction

Flow cytometrically sorted cells were resuspended at 1×10^6^ cells/mL and loaded on Chromium Controller (10X Genomics) to obtain single cells. For scRNA-seq library construction, Chromium Single Cell 3’ Library and Gel Bead Kit V2 (10X Genomics, PN120237) was used to generate single cell gel beads in emulsion (GEM). The captured cells were lysed, and the released RNA was reverse-transcribed with primers containing poly-(d) T, barcode, Unique Molecular Identifiers (UMIs) and read 1 primer sequence in GEMs. Barcoded cDNA was purified and amplified by PCR. The adapter ligation reaction was performed to add sample index and read 2 primer sequence. After quality control, the libraries were sequenced on Illumina Novaseq 6000 platform in 150 bp pair-end manner.

### Processing of scRNA-seq data

Sequencing data from 10X Genomics were processed with CellRanger (version 3.0.1) for demultiplexing, barcode processing and single-cell 3’ gene counting. Mouse (mm10) or human (GRCh37) reference genome was used for sequence alignment. Only confidently mapped, non-PCR duplicates with valid barcodes and UMIs were used to generate the gene-barcode matrix. For E12.5 mouse mandibles, cells with more than 2000 reads and less than 3% of mitochondrial genes were retained for downstream analyses. For adult mouse mandibles, cells with more than 200 reads and less than 10% of mitochondrial genes were retained for downstream analyses. For human mandible data, cells with more than 5000 reads and less than 10% of mitochondrial genes were retained for downstream analyses. Cell doublets were removed using Doublet Finder software implemented in R. Clustering and visualization were performed by Seurat. *LogNormalize*, a global-scaling normalization method, was employed to normalize gene expression. The expression measurement of a transcript in one cell was divided by all transcripts of the cell and multiplied by a scale factor (10,000 by default), followed by logarithmic transformation. *FindVariableFeatures* was used to get top 2,000 variable features per dataset, and a linear transformation (*ScaleData*) was applied to scale the data.

### Dimensionality reduction, clustering, and visualization

Principal component analysis (PCA) of the scaled data was performed. Top 20 principal components for E12.5 mandible data, top 12 principal components for adult mandible data, and top 11 principal components for human mandible data were used in the following steps. UMAP dimensional reduction technique was used to visualize and explore these datasets. *FindClusters* with resolution of 0.21 (for E12.5 mandible data), 0.23 (for adult mouse mandible data), or 0.2 (for human mandible data) was used to identify clusters for each dataset. *FindAllMarkers* with the default parameters (except that “logfc.threshold=0.25”) was used to find markers for different cell clusters. All markers were filtered with “p_adj_val” less than 0.05

### Cell-cell interaction analysis

Cell-cell interaction analysis was performed by the CellChat package embedded in R. The expression matrix and information of cell types were inputted to construct the CellChat object. Database subset “Secreted Signaling” of ligand and receptor pairs was used (CellChat.mouse). The communication among clusters was calculated by *computeCommunProb* and *computeCommunProbPathway* function. The contribution of pathways in different clusters was shown by network plot.

### Regulon analysis

Regulon analysis was performed using the SCENIC package implemented in python. The initial co-expression gene regulatory networks (GRN) were built by *grn* function with grnboost2 method. TF motifs were generated by *ctx* function using the mm10-tss-centered-10kb (for mouse) or hg19-tss-centered-10kb (for human) database. The AUC (area under the curve) score was calculated by *aucell* function. The mean regulon activity scores for each cluster were calculated and the top 20 regulons were visualized by heatmap. Binary activity of regulon was calculated by *binarize* function (binarization package implemented in python).

### Pseudotime analysis

Pseudotime analysis was performed using the Monocle3 package, which infer trajectory by fitting differentiation-principal curves based on given cell embeddings. Cds object was constructed by *new_cell_data_set* function based on expressional matrix, cell type and gene annotation information calculated by Seurat. The *learn_graph* function was used to infer trajectory based on Seurat UMAP coordinates. By specifying the starting cluster, cells were assigned onto the differentiation trajectory, and *order_cells* function was used to order genes for trajectory reconstruction. The mdSSC population was set as the starting cluster for both E12.5 and adult mouse mandible data.

### GO and KEGG analysis

Differentially expressed genes (DEGs) in each cluster were used to perform GO and KEGG pathway enrichment analysis by clusterProfiler package. The significantly expressed genes (marker genes) of each cluster were used as input and ontology was set to BP (biological process). The enriched GO terms were filtered by setting *P-value* cutoff to 0.05. Simplified function was performed to select the most significantly enriched terms. Dotplot was drawn by ggplot2, with “GeneRatio” as horizontal axis and “Term” as longitudinal axis.

### Bulk RNA-seq library construction and data analysis

Foxp4^+^ SSCs and Foxp4^-^ SSCs were isolated by flow cytometry as described above. mRNA was purified from total RNA using poly-T beads, fragmented, and reverse-transcribed with random hexamers. Second-strand synthesis used dUTP instead of dTTP to preserve strand information. Directional libraries were prepared by end repair, A-tailing, adapter ligation, size selection, USER digestion, amplification, and purification. Library quality was assessed by Qubit, qPCR, and Bioanalyzer. Qualified libraries were pooled and sequenced on an Illumina platform. Raw reads (FASTQ) were processed with fastp to remove adapter, poly-N, and low-quality reads. Reads were discarded in case of adapter contamination, >10% uncertain bases, or >50% bases with Phred quality≤5. Q20, Q30, and GC content were then calculated. Clean reads were used for downstream analyses. Gene expression was quantified as CPM (counts per million) for PCA and Pearson correlation analyses. DEGs were calculated by DESeq2 (|log2FC| > 0.25, padj < 0.1). GO and KEGG enrichment analyses were performed as described above.

## QUANTIFICATION AND STATISTICAL ANALYSIS

Statistical analyses were performed using GraphPad Prism 8. Detailed statistical methods were specified in the figure legends. Comparisons between two groups were analyzed using unpaired two-tailed Student’s t test. Comparisons between multiple groups were analyzed using one-way ANOVA followed by Tukey’s post hoc test. All data represent mean ± SD, \**P* < 0.05, \*\**P* < 0.01, \*\*\**P* < 0.001.

---

**Figure S1.**
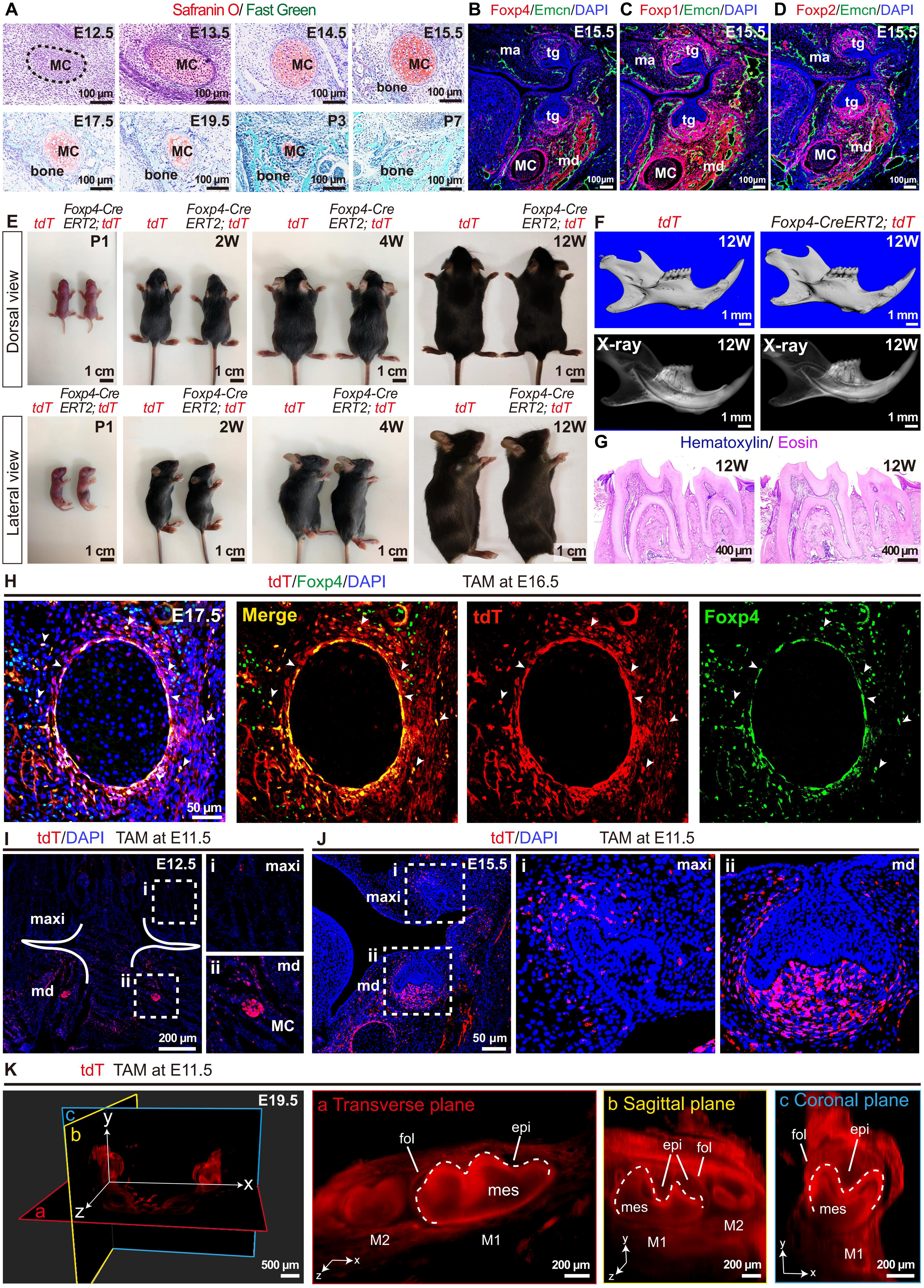
Characterization of MC development, *Foxp4-CreERT2* mice, and distribution of Foxp4^+^ cells in the maxilla and mandible. Related to Figure 1. (A) Safranin O/Fast Green staining of E12.5 to P7 mandibles in wild-type mice. Dotted circle indicated that mesenchymal condensation that gives rise to Meckel’s cartilage (MC) at E12.5. (B-D) IF images showing Foxp4 (B), Foxp1 (C), Foxp2 (D) expression in E15.5 wild-type embryos. MC: Meckel’s cartilage, md: mandible, ma: maxilla, tg: tooth germ. (E) The dorsal (upper) and lateral (lower) views of *Rosa26-tdTomato* (left) and *Foxp4-CreER; tdTomato* (right) mice from P1 to 12 weeks old. (F) MicroCT images showing comparable mandible and tooth development in *Rosa26-tdTomato* and *Foxp4-CreERT2; tdTomato* mice at 12 weeks old. (G) Hematoxylin/Eosin images showing comparable tooth development in *Rosa26-tdTomato* and *Foxp4-CreERT2; tdTomato* mice at 12 weeks old. (H) IF images of E17.5 mandibles in *Foxp4-CreERT2; tdTomato* mice. Tamoxifen was administered for 1 day at E16.5. The endogenous Foxp4 antibody was for the immunostaining. The tdTomato (tdT), Foxp4 and DAPI signals were shown (n=3 mice from 3 independent experiments). (I and J) Fluorescence images of E12.5 and E15.5 mandibles in *Foxp4-CreERT2; tdTomato* mice. Tamoxifen was administered for 1 day at E11.5. The tdTomato (tdT) and DAPI signals were shown. The solid curve indicated the border between maxilla (maxi) and mandible (md). Absence of tdTomato signal in E12.5 maxilla (i) and evident tdTomato signal indicating MC in E12.5 mandible (ii) were enlarged (I). Developing tooth germs in the maxilla (i) and mandible (ii) were enlarged (J) (n=3 mice from 3 independent experiments). (K) Light-sheet IF image of E19.5 mandible in *Foxp4-CreERT2; tdTomato* mice. Tamoxifen was administered for 1 day at E11.5. The tdTomato (tdT) signal on transverse (a), sagittal (b) and coronal section planes (c) were shown. M1: first molar; M2: second molar. fol: dental follicle, epi: dental epithelium, mes: dental mesenchyme.

**Figure S2.**
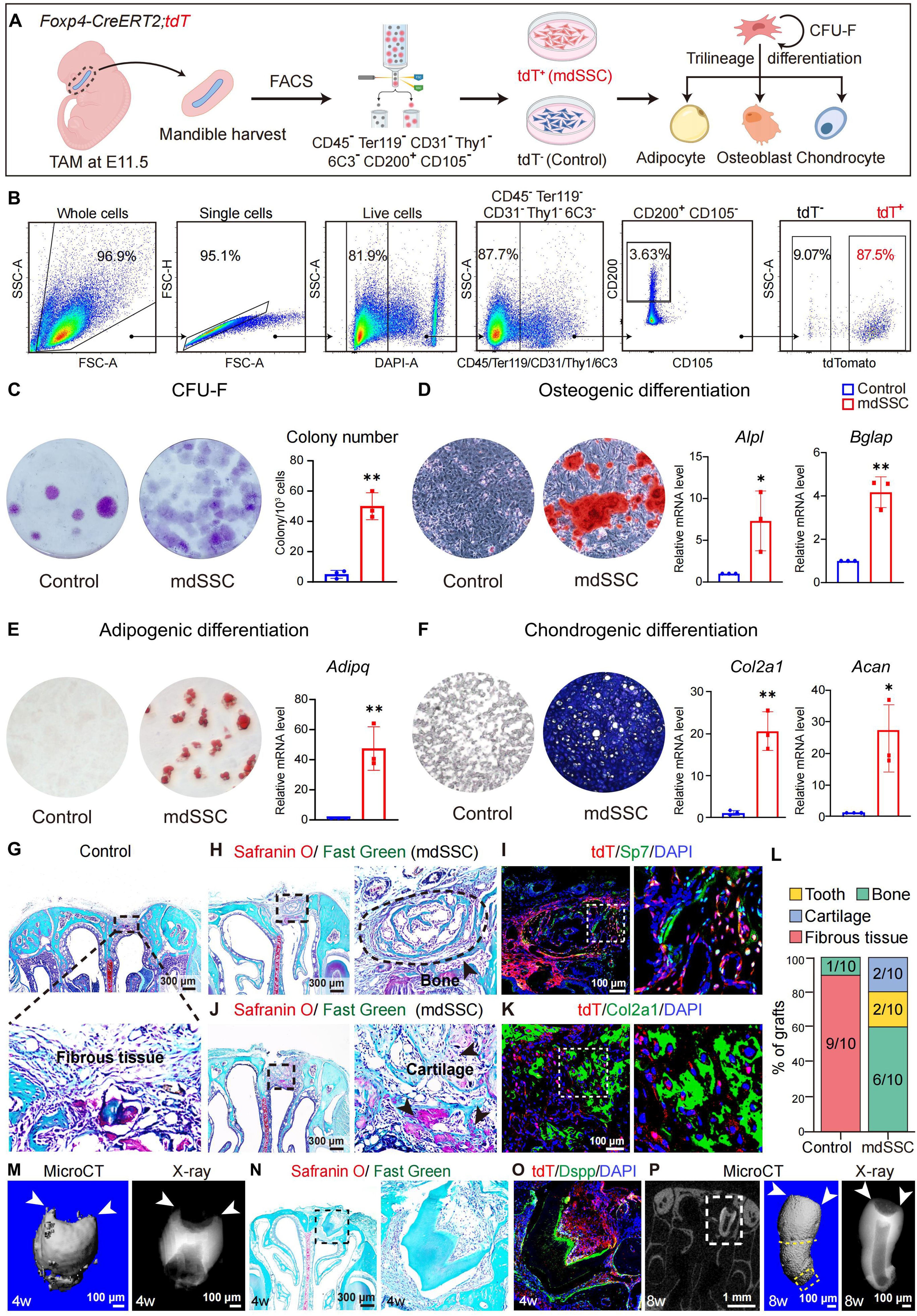
Gating strategies and functional characterization of Foxp4^+^ mdSSCs from E12.5-E16.5 mouse embryos. Related to Figure 2. (A) Sampling workflow and experimental scheme. Tamoxifen was administered for 1 day at E11.5. Cell sorting was performed at E12.5. (B) Gating strategy for sorting mdSSCs and control cells. (C) Crystal violet staining of CFU-F colonies with quantification (n=3 independent experiments). (D-F) *In vitro* tri-lineage differentiation. Alizarin red (D), Oil Red O (E) and Toluidine blue (F) staining showing osteogenic, adipogenic and chondrogenic differentiation, respectively. qPCR analyses of adipogenic, osteogenic and chondrogenic markers were shown (n=3 independent experiments). (G) Safranin O/Fast Green staining of the fibrous tissue formed by Foxp4^-^ SSCs (Control) 4 weeks after nasal sinus transplantation. (H and J) Safranin O/Fast Green staining of the ectopic bone (H) and cartilage (J) formed by mdSSCs 4 weeks after nasal sinus transplantation. Dotted circle indicated the ectopic bones. Arrowheads indicated the ectopic cartilage. (I and K) IF images of tdTomato (tdT), Sp7 (I), Col2a1 (K) and DAPI in the ectopic bone and cartilage formed by Foxp4^+^ mdSSCs 4 weeks after nasal sinus transplantation. (L) Quantification of the 4-week nasal sinus transplantation data (n=10 mice per group from 6 independent experiments). (M and N) Representative microCT images (M) and Safranin O/Fast Green staining (N) of the ectopic tooth formed by E15.5 mdSSCs 4 weeks after nasal sinus transplantation. (O) IF images of tdTomato (tdT) (red), Dspp (green) and DAPI (blue) in the ectopic tooth (N). (P) Representative microCT images of the ectopic tooth formed by E16.5 mdSSCs 8 weeks after nasal sinus transplantation. Snapshot (left), 3D (middle) and 2D views (right) were shown. Statistical significance was determined by unpaired two-tailed Student’s t test. All data represent mean ± SD (∗p < 0.05, ∗∗p < 0.01).

**Figure S3.**
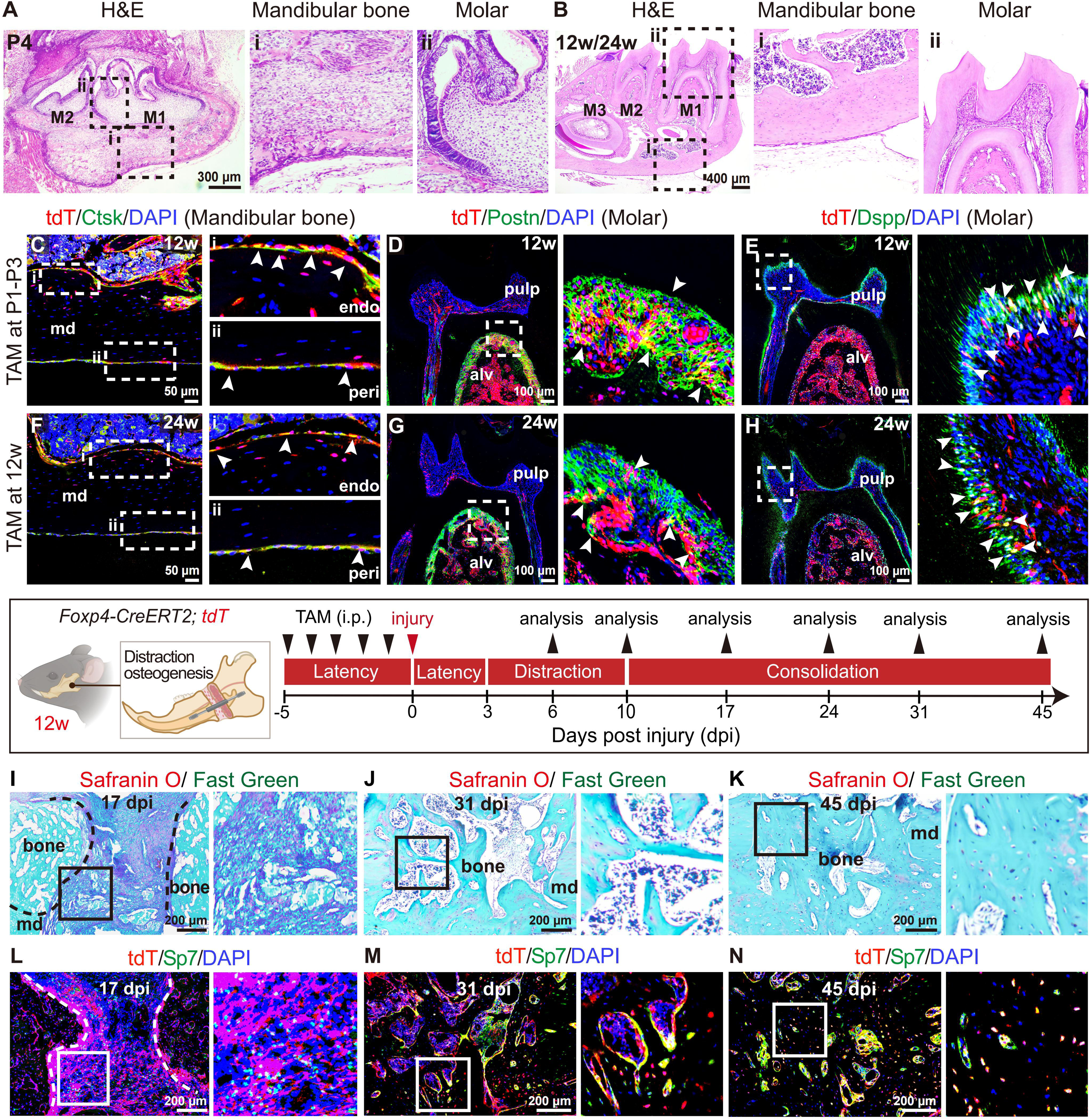
Lineage tracing of Foxp4^+^ cells in postnatal mandible under steady state or during fracture repair. Related to Figure 3. (A and B) Representative H&E staining of P4 (A) and 12-week-old (B, same imaging positions for 24-week-old mice) mandible sections. The mandibular bone beneath the M1 molar roots (i) and M1 molar (ii) were enlarged. (C-E) IF images of tdTomato (tdT), Ctsk, Postn, Dspp and DAPI in mandible sections of 12-week-old *Foxp4-CreERT2; tdTomato* mice. Tamoxifen was administered for 3 consecutive days at P1-P3. The endosteal (endo, i) and periosteal (peri, ii) regions of the mandibular bones (C), the periodontal ligament (D), as well as the dental pulp of the molar (E) were enlarged. Arrowheads indicated tdT^+^ endosteal/periosteal cells (C), periodontal ligament cells (D) and odontoblasts cells (E). md: mandibular bone, alv: alveolar bone. (F-H) IF images of tdTomato (tdT), Ctsk, Postn, Dspp and DAPI in mandible sections of 24-week-old *Foxp4-CreERT2; tdTomato* mice. Tamoxifen was administered for 5 consecutive days at 12-week-old. The endosteal (endo, i), periosteal (peri, ii) regions of the mandibular bones (F), the periodontal ligament regions (G) as well as the dental pulp of the molar (H) were enlarged. Arrowheads indicated tdT^+^ endosteal/periosteal cells (F), periodontal ligament cells (G) and odontoblasts cells (H). md: mandibular bone, alv: alveolar bone. (I-K) Safranin O/Fast Green staining of mandibular bone repair. DO-operated *Foxp4-CreERT2; tdTomato* mice were analyzed at 17, 31, and 45 dpi. Dotted lines indicated the osteogenic front. md: mandible, bone: newly formed woven bone. (L-N) IF images of 17, 31, and 45 dpi mandibular bones in *Foxp4-CreERT2; tdTomato* mice. The tdTomato (tdT), Sp7 and DAPI signals were shown.

**Figure S4.**
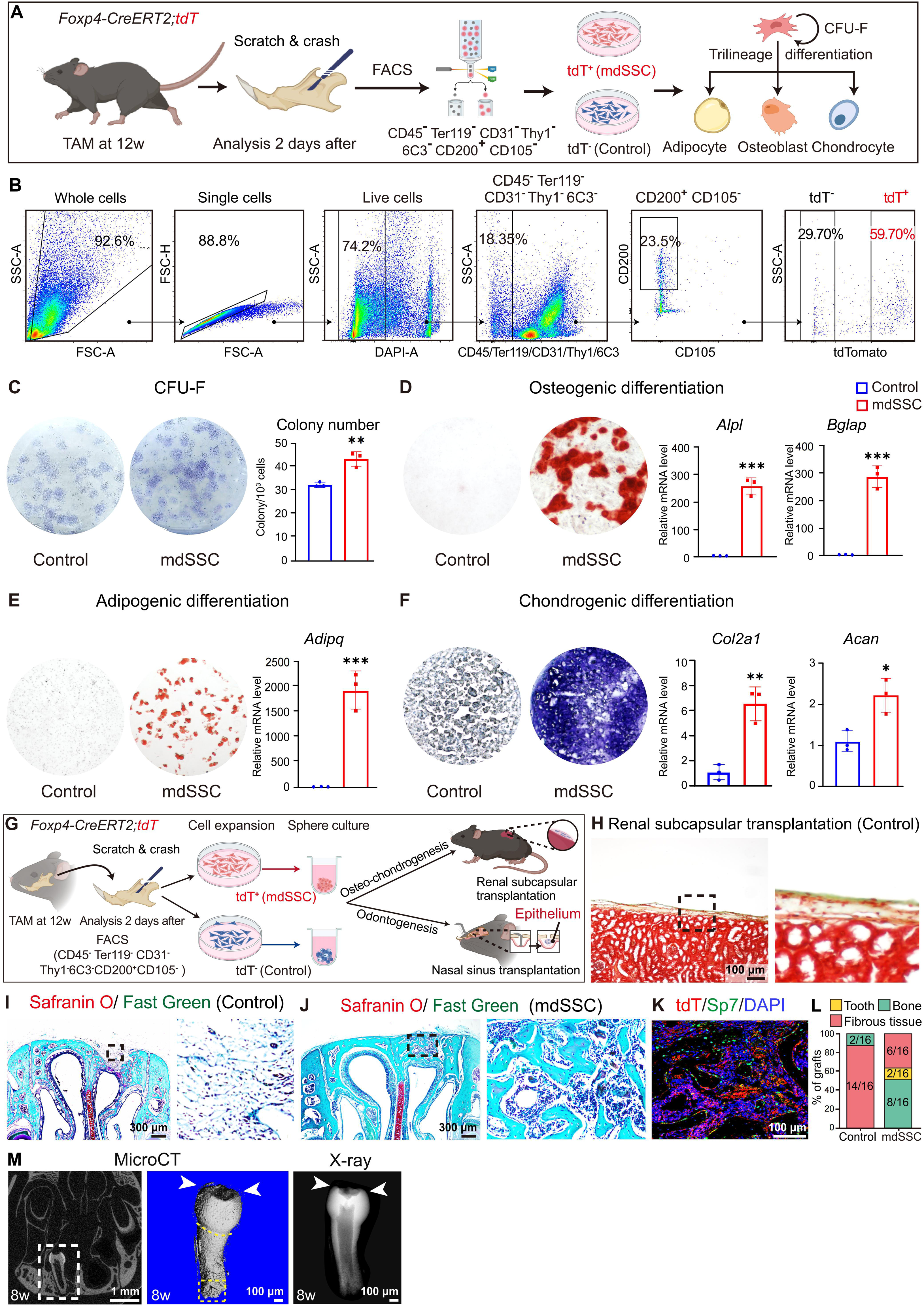
Gating strategies and the functional characterization of Foxp4+ mdSSCs from adult mice. Related to Figure 4. (A) Sampling workflow and experimental scheme. Cells were sorted 2 days after tamoxifen administration (5 consecutive days starting from 12 weeks of age). (B) Gating strategy for sorting mdSSC and control cells. (C) Crystal violet staining of CFU-F colonies with quantification (n=3 independent experiments). (D-F) In vitro tri-lineage differentiation. Alizarin red (D), Oil Red O (E) and Toluidine blue (F) staining showing osteogenic, adipogenic and chondrogenic differentiation, respectively. qPCR analyses of adipogenic, osteogenic and chondrogenic markers were shown (n=3 independent experiments). (G) Transplantation workflow and experimental scheme. Cells were sorted 2 days after tamoxifen administration (5 consecutive days starting from 12 weeks of age). (H) Movat’s pentachrome staining showing fibrous tissue formation 4 weeks after renal subcapsular transplantation of Foxp4- SSCs (Control). (I) Safranin O/Fast Green staining of the fibrous tissue formed by Foxp4- SSCs (Control) 4 weeks after nasal sinus transplantation. (J) Safranin O/Fast Green staining of the ectopic bone formed by adult mdSSCs 4 weeks after nasal sinus transplantation. (K) IF images of tdTomato (tdT), Sp7 and DAPI in the ectopic bone formed by adult mdSSCs 4 weeks after nasal sinus transplantation. (L) Quantification of the 4-week nasal sinus transplantation data (n=16 mice per group from 5 independent experiments). (M) Representative microCT images of the ectopic tooth formed by adult mdSSCs 8 weeks after nasal sinus transplantation. Snapshot (left), 3D (middle) and 2D views (right) were shown. Statistical significance was determined by unpaired two-tailed Student’s t test. All data represent mean ± SD (∗p < 0.05, ∗∗p < 0.01, ∗∗∗p < 0.001).

**Figure S5.**
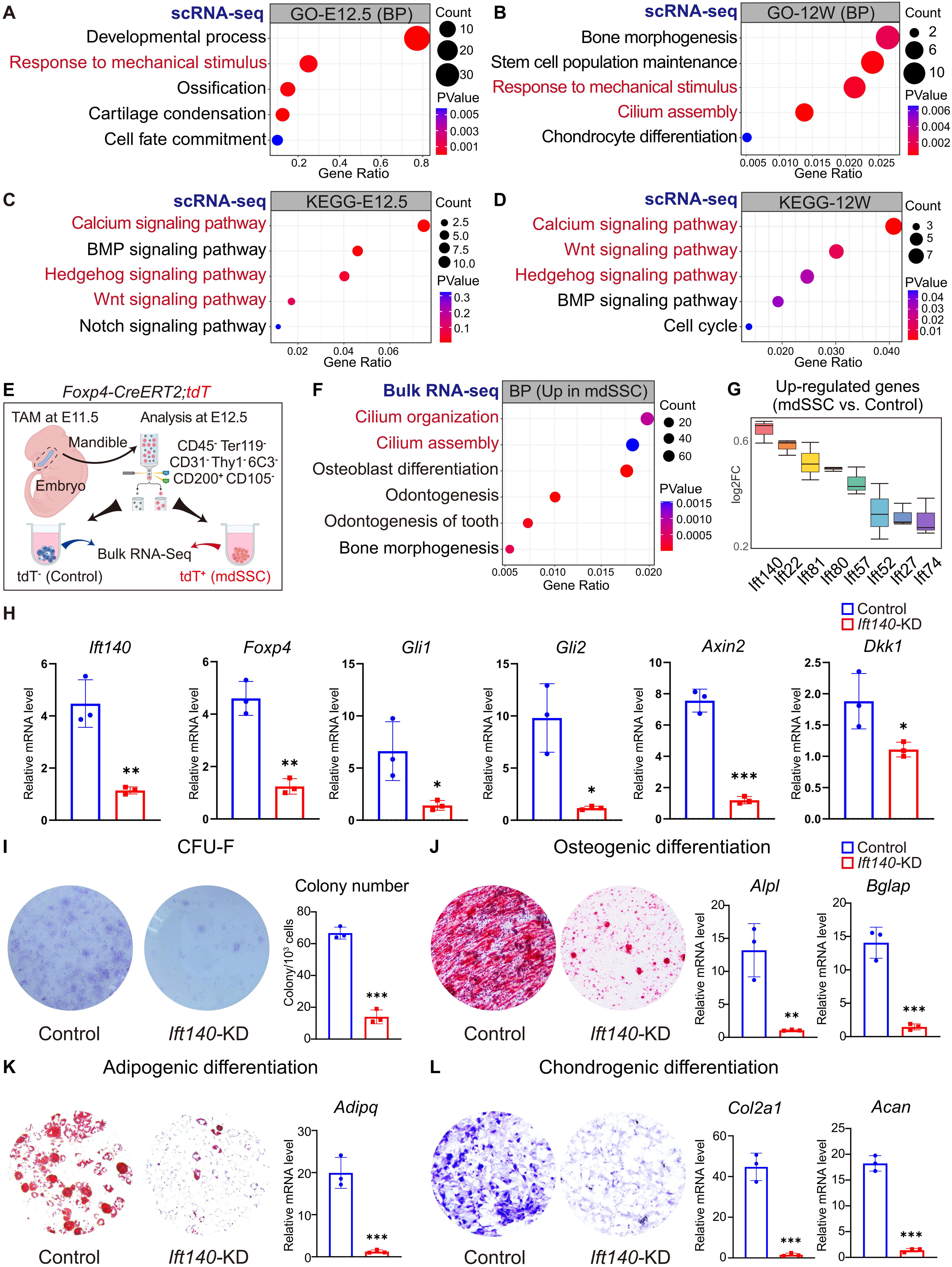
Bioinformatic and functional analysis of primary cilia and Ift140 in mdSSCs. Related to Figure 6. (A-D) Dot plots showing GO terms and KEGG pathways enriched in the mdSSC subset of E12.5 and 12-week-old mandible scRNA-seq data. BP: Biological process. (E) Bulk RNA-seq workflow and experimental scheme. Tamoxifen was administered for 1 day at E11.5. Mouse mandibles were analyzed at E12.5. (F) Dot plot showing GO terms enriched in Foxp4+ SSCs (mdSSC) as compared to Foxp4-SSCs by analyzing E12.5 bulk RNA-seq data (E). BP: Biological process. (G) Box plot showing relative enrichment of intraflagellar transport genes in Foxp4+ SSCs (mdSSC) as compared to Foxp4- SSCs by analyzing E12.5 bulk RNA-seq data (E). (H) qPCR analysis of Foxp4+ mdSSCs following Ift140 or control siRNA knockdown (n=3 independent experiments). (I) Crystal violet staining of CFU-F colonies with quantification (n=3 independent experiments). (J-L) In vitro trilineage differentiation. Alizarin red (J), Oil Red O (K) and Toluidine blue (L) staining showing osteogenic, adipogenic and chondrogenic differentiation, respectively. qPCR analyses of adipogenic, osteogenic and chondrogenic markers were shown (n=3 independent experiments). Statistical significance was determined by unpaired two-tailed Student’s t test. All data represent mean ± SD (∗p < 0.05, ∗∗p < 0.01, ∗∗∗p < 0.001).

**Figure S6.**
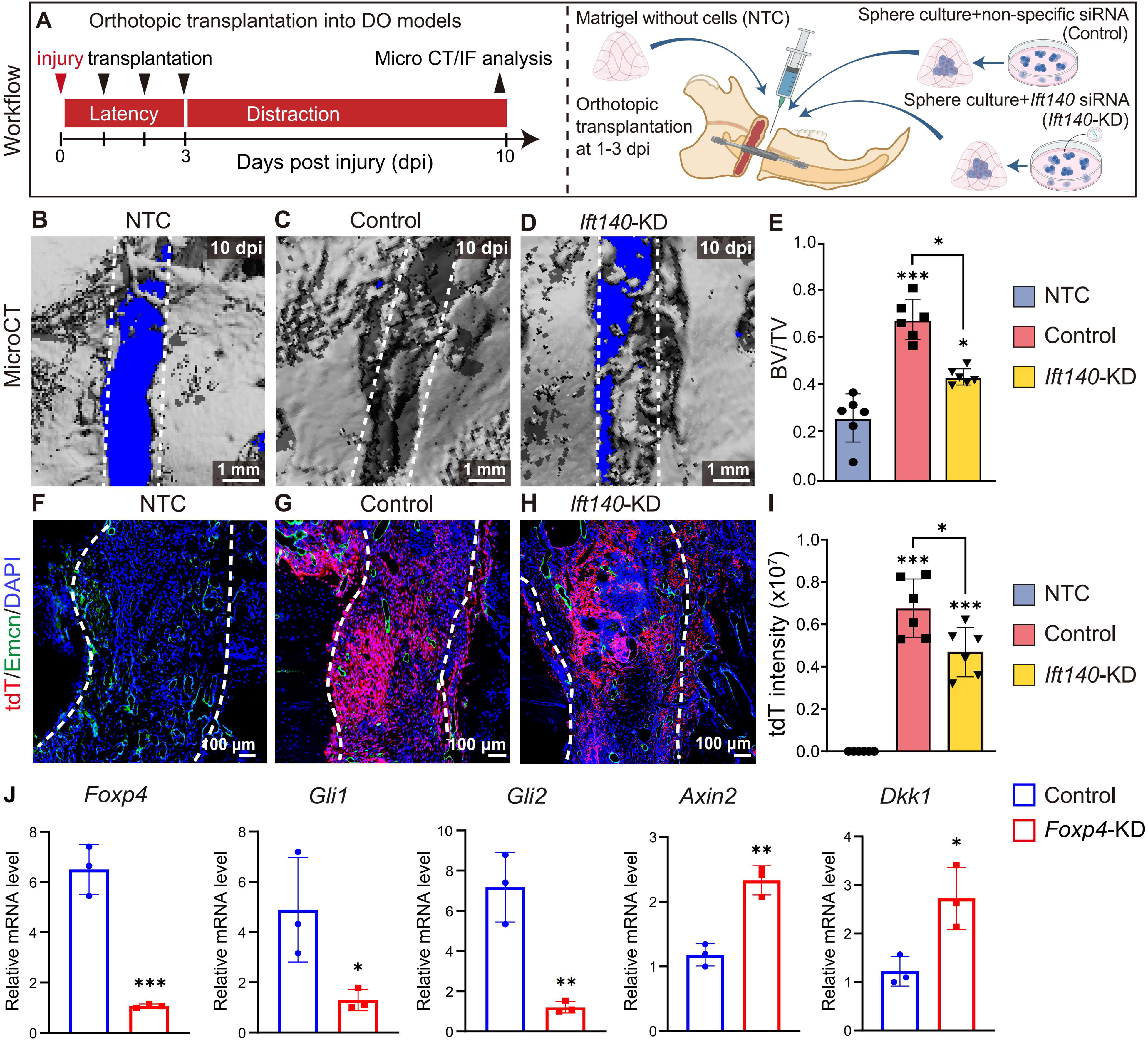
Ift140 and Foxp4 tightly regulate mdSSC function. Related to Figure 6. (A) Experimental scheme of mdSSC orthotopic transplantation into wild-type mice after DO. Negative control (NTC, Matrigel only), control siRNA or Ift140 siRNA knockdown E12.5 mdSSCs were transplanted into the mandibular fracture site within the first three days and analyzed at 10 dpi. (B-E) Representative microCT images of mandible repair at 10 dpi (B-D) with quantification of BV/TV at the fracture site (E). Dotted lines indicated the regions analyzed for new bone formation (n=6 mice per group from three independent experiments). (F-I) IF images of mandible repair at 10 dpi (F-H) with quantification of tdTomato (tdT) signal intensity (I). Dotted lines indicated the osteogenic front (n=6 mice per genotype from 3 independent experiments). (J) qPCR analysis of Foxp4+ mdSSCs following Foxp4 or control siRNA knockdown (n=3 independent experiments). Statistical significance was determined by one-way ANOVA followed by Tukey’s post hoc test (E and I), or unpaired two-tailed Student’s t test (J). All data represent mean ± SD (∗p < 0.05, ∗∗p < 0.01, ∗∗∗p < 0.001).

**Figure S7.**
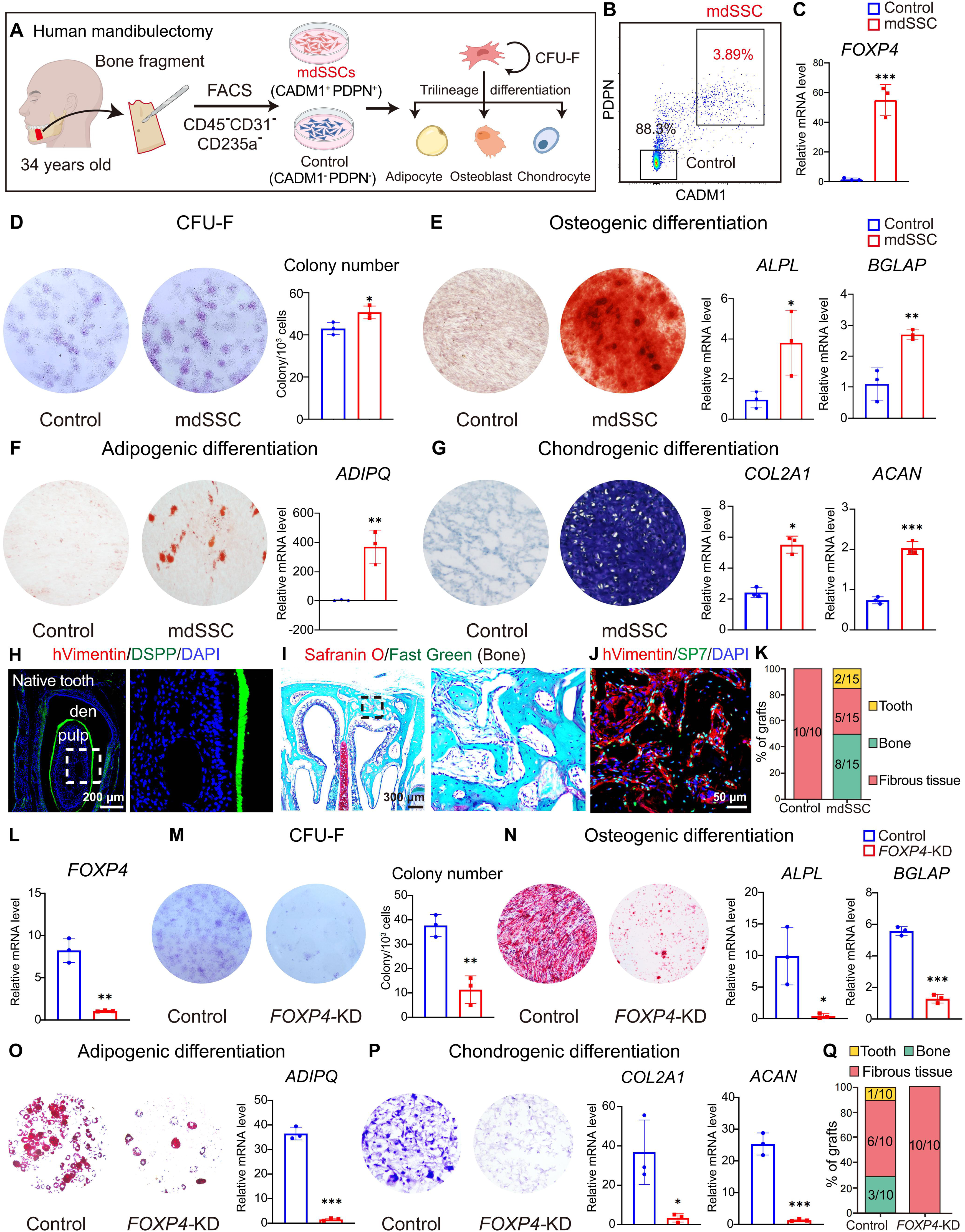
Gating strategies and the functional characterization of human FOXP4+ mdSSCs. Related to Figure 7. (A) Sampling workflow and experimental scheme. The mandibular bone fragment obtained from a 34-year-old patient was analyzed. (B) Gating strategy for sorting human mdSSC and control cells. (C) qPCR analysis of FOXP4 expression in human mdSSC and control cells (n=3 independent experiments). (D) Crystal violet staining of CFU-F colonies with quantification (n=3 independent experiments). (E-G) In vitro tri-lineage differentiation. Alizarin red (E), Oil Red O (F) and Toluidine blue (G) staining showing osteogenic, adipogenic and chondrogenic differentiation, respectively. qPCR analyses of adipogenic, osteogenic and chondrogenic markers were shown (n=3 independent experiments). (H) IF images of human Vimentin (hVimentin), DSPP and DAPI in the native tooth of recipient mice (NCG). den: dentin. (I) Safranin O/Fast Green staining of the ectopic bone formed by human mdSSCs after nasal sinus transplantation into NCG immunodeficient mice. (J) IF images of human Vimentin (hVimentin), SP7 and DAPI in the ectopic bone formed by human mdSSCs after nasal sinus transplantation into NCG immunodeficient mice. (K) Quantification of nasal sinus transplantation of human mdSSCs (n=10 or 15 recipient mice per group from 5 independent experiments). (L) qPCR analysis in human mdSSCs following FOXP4 or control siRNA knockdown (n=3 independent experiments). (M) Crystal violet staining of CFU-F colonies with quantification (n=3 independent experiments). (N-P) In vitro trilineage differentiation. Alizarin red (N), Oil Red O (O) and Toluidine blue (P) staining showing osteogenic, adipogenic and chondrogenic differentiation, respectively. qPCR analyses of adipogenic, osteogenic and chondrogenic markers were shown (n=3 independent experiments). (Q) Quantification of the 4-week nasal sinus transplantation data for human mdSSCs following FOXP4 or control siRNA knockdown (n=10 recipient mice per group from 3 independent experiments). Statistical significance was determined by unpaired two-tailed Student’s t test. All data represent mean ± SD (∗p < 0.05, ∗∗p < 0.01, ∗∗∗p < 0.001).

