## Supplementary material for "Foxp4^+^ Mandibular Skeletal Stem Cells Orchestrate Bone/Tooth Development and Regeneration": Table S1

| Genotyping PCR primer list | | | | |
|---|---|---|---|---|
| **Strains** | **Primer 1** | **Primer 2** | **Primer 3** | **Primer 4** |
| Ift140$^{fl/fl}$ | ATCTTAATTTGTGTTGAAGGGGTT | CTGCCAGGGGTACATGGTAGTAAG | NA | NA |
| Foxp4-CreERT2 | AGCCCCTCCCCCATGTCTCTTCTC | GGCGCCTGTCGGTCCTTATGG | GCAAGGGCCGTTTACTGGTG | GCCTGGC GATCCCTGAACA |
| Rosa26$^{tdTomato}$ | AAGGGAGCTGCAGTGGAGTA | CTGTTCCTGTACGGCATGG | CCGAAAATCTGTGGGAAGTC | GGCATTAAAGCAGCGTATCC |
| Rosa26$^{DTA}$ | GTTATCAGTAAGGGAGCTGCAGTGG | GGCGGATCACAAGCAATAATAACC | AAGACCGCGAAGAGTTTGTCCTC | NA |

| qPCR primer list (Human) | | |
|---|---|---|
| **Gene** | **Forward primer** | **Reverse primer** |
| GAPDH | GACCCCTTCATTGACCTCAAC | CTTCTCCATGGTGGTGAAGA |
| FOXP4 | GTTCACCAGGATGTTCGCCT | CTCCGCTTCTGATACTCCCG |
| ADIPOQ | GCTCTGTGCTCCTGCATCT | TTACGCTCTCCTTCCCCATA |
| COL2A1 | GTCCTGTCGGTCCTGCTG | GAGGACCTTGAGCACCTTCA |
| ACAN | ACTCTGGGTTTTCGTGACTCT | ACACTCAGCGAGTTGTCATGG |
| ALPL | GCCTACACGGTCCTCCTATACGG | CACTGCTGACTGCTGCCGATAC |
| BGLAP | CACTCCTCGCCCTATTGGC | CCCTCCTGCTTGGACACAAAG |
| AXIN2 | TAACCCCTCAGAGCGATGGA | AGTTCCTCTCAGCAATCGGC |
| DKK1 | ACGCTATCAAGAACCTGCCC | GGGTACGGCTGGTAGTTGTC |
| GLI1 | AGACAGAGGCCCACTCTTTTC | GCCAATGGAGAGATGACCGT |
| GLI2 | CCTGCGTGCTAGAGGCAAAC | TCGCATGTCAATCGGTAGGG |
| IFT140 | AGTGTGGAAGCACGTCTGAG | AATAGCCGAAGGTGAGCGAG |

| qPCR primer list (Mouse) | | |
|---|---|---|
| **Gene** | **Forward primer** | **Reverse primer** |
| β-Actin | GCTCTTTTCCAGCCTTCCTT | CTTCTGCATCCTGTCAGCAA |
| Adipoq | TGTTCCTCTTAATCCTGCCCA | CCAACCTGCACAAGTTCCCTT |
| Col2a1 | GTCCCCCTGGCCTTAGTG | CCACCAGCCTTCTCGTCA |
| Acan | TGAAGCAGAAGGTCTGGACA | CCAGAAGGAATCCCACTAACA |
| Alpl | CGGATCCTGACCAAAAACC | TCATGATGTCCGTGGTCA AT |
| Bglap | CTGACCTCACAGATGCCAAGC | TGGTCTGATAGCTCGTCACAAG |
| Axin2 | CGCCTAGTGACTGCTGGAAA | ACGGAAAACAACGATCCCGA |
| Dkk1 | TCCAACGCGATCAAGAACCT | GTACTTGTTCCCGCCCTCAT |
| Gli1 | CTTCCCTGAGACGCCATGTT | ACCCTGGGACCCTGACATAA |
| Gli2 | CTCTGGCTCTTGTGGTGTGG | GTCAAGGGAAAAGCAGATCAACA |
| Foxp4 | CAACCTCAGCCTGCACAAGTGTTT | TCTGCAGGCTCCTCTTTGACTTGT |
| Ift140 | GTAGGCAGGATGAACTCGGG | CCAGTTTGGGTTTTCCGCTG |

| siRNA primer list | | | |
|---|---|---|---|
| **Gene** | **Species/Gene ID** | **Sense（5'-3'）** | **Antisense（5'-3'）** |
| Ift140 | mouse/106633 | GCAUCUAUAAGCCCAUCUU | AAGAUGGGCUUAUAGAUGC |
| Sox9 | mouse/20682 | CGACGUGGACAUCGGUGAA(dT)(dT) | UUCACCGAUGUCCACGUCG(dT)(dT) |
| Msx2 | mouse/17702 | GCAAGAGGCGGAACUGGAA(dT)(dT) | UUCCAGUUCCGCCUCUUGC(dT)(dT) |
| Lhx8 | mouse/16875 | GCAAGACGUCAAUCAUCCA(dT)(dT) | UGGAUGAUUGACGUCUUGC(dT)(dT) |
| Meis1 | mouse/17268 | CCAUGAUAGACCAGUCCAA(dT)(dT) | UUGGACUGGUCUAUCAUGG(dT)(dT) |
| Foxp4 | mouse/74123 | CCACCAGAUACAAGUCAAA(dT)(dT) | UUUGACUUGUAUCUGGUGG(dT)(dT) |
| FOXP4 | human/116113 | UCGGCUCCAUCUGGUCAGAAU(dT)(dT) | AUUCUGACCAGAUGGAGCCGA(dT)(dT) |
